# Mechanism of praziquantel action at a parasitic flatworm ion channel

**DOI:** 10.1101/2021.03.09.434291

**Authors:** Sang-Kyu Park, Lukas Friedrich, Nawal A. Yahya, Claudia Rohr, Evgeny G. Chulkov, David Maillard, Friedrich Rippmann, Thomas Spangenberg, Jonathan S. Marchant

## Abstract

Praziquantel (PZQ) is an essential medicine for treating parasitic flatworm infections such as schistosomiasis, which afflicts over 250 million people. However, PZQ is not universally effective, lacking activity against the liver fluke *Fasciola*. The reason for this insensitivity is unclear, as the mechanism of PZQ action is unknown. Here, we show PZQ activates a transient receptor potential melastatin ion channel (TRPM_PZQ_) in schistosomes by engaging a hydrophobic ligand binding pocket within the voltage-sensor like domain to cause Ca^2+^ entry and worm paralysis. PZQ activates TRPM_PZQ_ homologues in other PZQ-sensitive flukes, but not *Fasciola*. However, a single amino acid change in the *Fasciola* TRPM_PZQ_ binding pocket, to mimic schistosome TRPM_PZQ_, confers PZQ sensitivity. After decades of clinical use, the basis of PZQ action at a druggable TRP channel is resolved.

---

Praziquantel (PZQ) is an essential medicine for treating diseases caused by parasitic flatworms (helminths). PZQ is the key therapy for schistosomiasis (Bilharzia), a disease of poverty that afflicts ~250 million people worldwide (*1, 2*). The use and continued efficacy of PZQ is integral to the WHO roadmap of interventions to control schistosomiasis morbidity and to eliminate schistosomiasis as a public health problem by 2030 (*3, 4*). WHO estimates that at least 290 million people required preventive treatment in 2018. PZQ is also employed to treat other fluke and cestode infections that cause human disease, but is ineffective against the common liver fluke *Fasciola. Fasciola* is a zoonotic pathogen that causes clinical and veterinary disease of livestock, responsible for considerable economic damage (*5*). The explanation for the varied effectiveness of PZQ against blood and liver flukes is unclear, as the mechanism of action of PZQ is unknown.

PZQ is administered as a racemic mixture with the (*R*)-PZQ enantiomer causing a rapid contraction of adult schistosome worms and damage to the worm surface while the (*S*)-PZQ enantiomer is considerably less effective (*6–9*). Recently, we identified a schistosome transient receptor potential (TRP) channel that is activated by PZQ, although like many previously proposed PZQ targets (*10, 11*), a direct binding site for PZQ on the target remains undefined (*12*). TRP channels are polymodal ion channels frequently employed in sensory signaling roles, responding to ligands and environmental cues. The *Schistosoma mansoni* TRP channel activated by PZQ (*Sm*.TRPM_PZQ_; Smp_246790.5, chromosome 3) belongs to the TRP melastatin subfamily, which in vertebrates contain members responsive to oxidative stress (TRPM2, (*13*)), high temperature (TRPM3, (*14*) as well as low temperature and cooling ligands such as menthol (TRPM8, (*15*)). Here, we coalesce ligand- and target-based approaches together with computational modelling to characterize how PZQ engages *Sm*.TRPM_PZQ_, with definition of a specific PZQ binding site revealing the molecular basis for the insensitivity of *Fasciola* to PZQ.

To characterize the pharmacological specificity of the schistosome TRP channel activated by PZQ (*Sm*.TRPM_PZQ_), we synthesized a series of 43 PZQ derivatives including several non-obvious analogues (Fig. S1, Table S1 and Supplementary Methods). The majority of analogues were structurally related to PZQ and the series included both resolved enantiomers (*R*)-PZQ (analogue **1**) and (*S*)-PZQ (analogue **2**), the major trans-(*R*)-4-OH PZQ metabolite (analogue **21**, (*9*)) and the veterinary drug epsiprantel (analogue **35**, Fig. 1A). Following transient expression of *Sm*.TRPM_PZQ_ in HEK293 cells, full concentration-response curves were performed for PZQ and each analogue, using a fluorescent assay to measure changes in intracellular Ca^2+^ (Fig. 1B and S2A). (*R*)-PZQ evoked a cytoplasmic Ca^2+^ signal with an EC_50_ of 0.46±0.12μM (Table S1). The enantiomer (*S*)-PZQ was ~50-fold less effective (EC_50_=24.7±1.3μM), consistent with the activity ratio of enantiomers seen *in vivo* (*9*). The entire EC_50_ dataset for these analogues at *Sm*.TRPM_PZQ_ (Fig. S2A, Table S1) was graphically represented as a color-coded heatmap (Fig. 1C). The majority of analogues displayed either no activity (19 compounds) or lower potency (13 compounds, EC_50_ values between 1-10μM; 7 compounds, EC_50_ values ≥10μM; Fig. 1C).

**Fig. 1.**
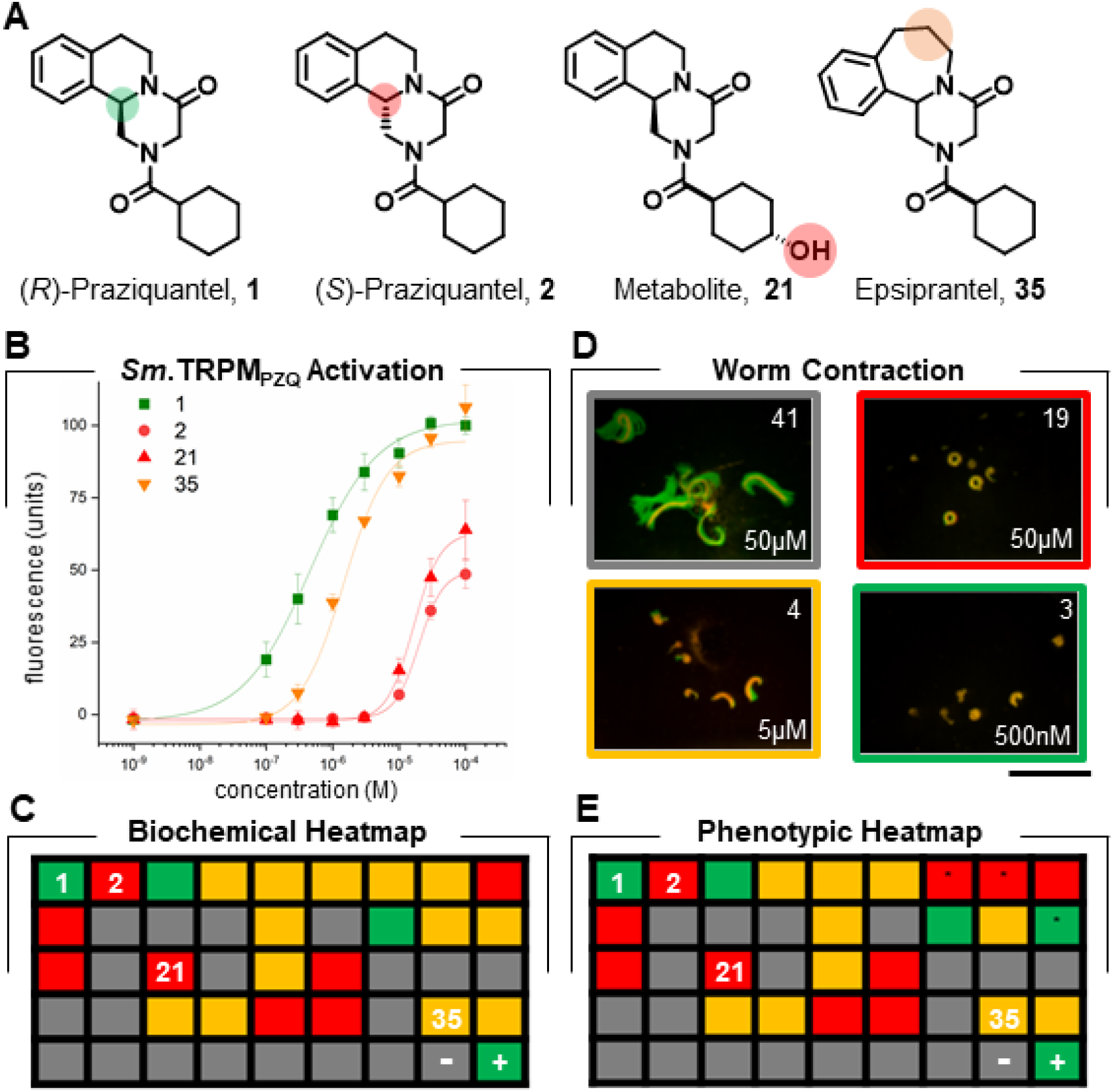
Structure-activity relationships of PZQ analogues. (**A**) Chemical structure of key compounds: (*R*)-PZQ (analogue #1), (*S*)-PZQ (analogue #2), the major metabolite (trans-(*R*)-4-OH PZQ, analogue #21), and epsiprantel (analogue #35) from the analogue collection fully indexed in Table S1. Colored circles highlight structural differences between these key analogues. (**B**) Concentration-response analysis for (*R*)-PZQ (analogue #1), (*S*)-PZQ (analogue #2), the major metabolite (trans-(*R*)-4-OH PZQ, analogue #21), and epsiprantel (analogue #35) on *Sm*.TRPM_PZQ_ assessed by peak fluo-4 fluorescence. Data from ≥3 independent experiments. (**C**) Graphical heatmap of the effects of each of the 43 analogues on *Sm*.TRPM_PZQ_ activation. Each analogue is numbered left to right (1 through 43) in the heatmap and indexed to Table S1, with responses to vehicle (-) and PZQ (+) indicated (bottom right). Colors represent EC_50_ values <1μM (green), 1-10μM (yellow), >10μM (red) or inactive (grey). (**D**) Images of adult schistosome worms with single frame image (red) overlaid with maximum intensity projection of a time-lapse series to illustrate effect of analogues on worm motion 5 mins after treatment with analogues 41 (top left), 19 (top right), 4 (bottom right) and 3 (bottom left) at indicated concentrations. Strong contractile responses are shown at different concentrations, or inactivity (analog 41), as coded in the heatmap in (E). Scalebar, 1cm. (**E**) Graphical heatmap of the effects of each of the 43 analogues on worm contraction ordered as per (C). Differences between the two heatmaps are asterisked.

As the relationship between target (*Sm*.TRPM_PZQ_) activation and worm phenotypic responses is unknown, we examined the ability of these same analogues to contract adult *S. mansoni* worms. Worm movement *ex vivo* was monitored before and after analogue addition. In the absence of drug exposure, movement of individual worms (red color) was evident by the area swept through by the worm in a maximal intensity projection of the recording (green color, Fig. 1D). The acute action of four different analogues (analogues **41, 19, 4** & **3**; Table S1) is shown in Figure 1D. These four analogues provide examples of dose-dependent inhibition of mobility over different concentrations ranges, or a lack of activity. An average motility score was calculated for each analogue relative to vehicle- and PZQ-treated worms (Fig. S2B) and used to produce a color-coded heatmap (Fig. 1E). Both the target (*Sm*.TRPM_PZQ_) and phenotypic (worm contraction) heatmaps showed similar structure-activity relationships (Fig. 1C and 1E), confirming the pharmacological fingerprint of *Sm*.TRPM_PZQ_ closely mirrors the activity of the same analogues at causing worm paralysis.

The structure-activity relationships (SAR) of these analogues revealed several trends. First, the cyclohexyl moiety in PZQ is a critical determinant for efficacy. Major modifications of this moiety yielded inactive or low potency analogs. Bulkier or more polar groups led to decreased potency with only minor alterations preserving comparable potency with (*R*)-PZQ (analogues **3** and **16**). Second, modifications of the three-ringed PZQ core (core structure ‘A’, Fig. S1) were poorly tolerated with exception of a fluorinated scaffold (core structure ‘B’, analogues **30** & **31**) or the enlarged piperidine ring of epsiprantel (Fig. 1A, analogues **35** & **36**). Third, stereochemically, all dextro- derivatives with the exception of (*S*)-PZQ were inactive. A ligand-based activity model confirmed the importance of these structural features (Fig. S2C), evidencing a ‘tight’ SAR with only small structural modifications of PZQ preserving activity at *Sm*.TRPM_PZQ_.

*Sm*.TRPMM_PZQ_ shows highest sequence homology to the ‘long’ TRPM channels, TRPM2 and TRPM8. The presence of a COOH terminal NUDT9H-enzyme like domain in *Sm*.TRPM_PZQ_ (Fig. 2A), diagnostic of TRPM2, may suggest a TRPM2-like activation mechanism. TRPM2 channels bind the channel activator ADP ribose at sites within the cytoplasmic regions in either the COOH-terminus (NUDT9H domain) and/or the NH2-terminus (TRPM homology region, MHR 1/2domain (*13, 16–19*). Truncation of the C-terminal NUDT9H domain of *Sm*.TRPM_PZQ_ did not impact responsivity to PZQ (Fig. 2A and S3A). *N*-terminal truncations into the MHR1/2 domains showed poor expression, consistent with the role of these domains in channel assembly and cell surface trafficking (*20*). However a panel of *N*-terminal point mutants targeting residues involved in ADP-ribose binding identified in other TRPM2 structures (*13, 17*) failed to prevent PZQ activation of *Sm*.TRPM_PZQ_ (Table S2).

**Fig. 2.**
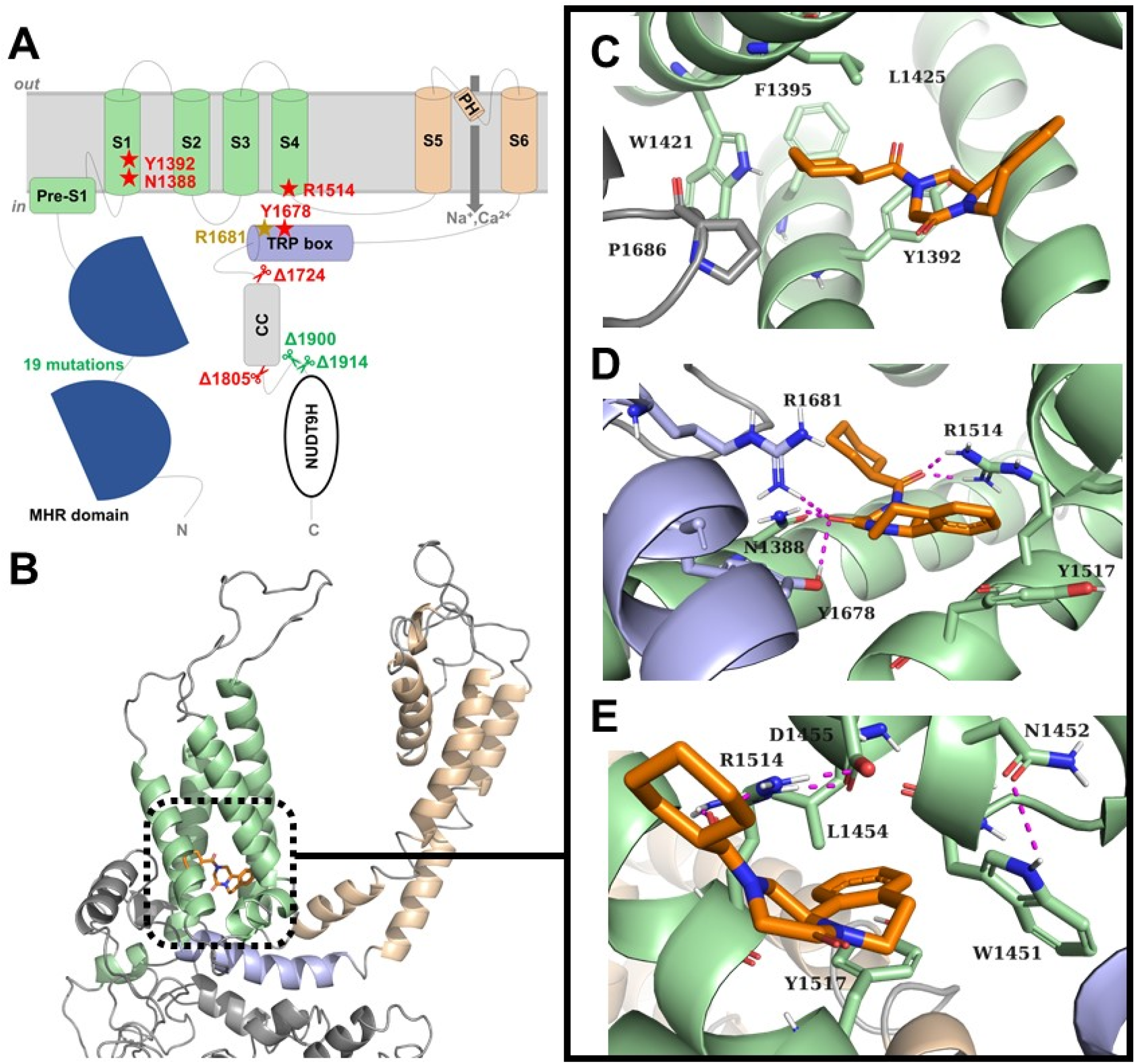
Modeling the PZQ binding pocket in *Sm*.TRPM_PZQ_. (**A**) Schematic of domain organization of *Sm*.TRPM_PZQ_ to highlight MHR domain, transmembrane spanning region (voltage-sensor-like domain, green; pore helices (PH), orange), TRP domain (blue), coiled-coil regions (CC) and COOH terminal NUDT9H domain. Truncation mutant (scissors) and specific point mutations (stars) analyzed in Figures S3A and S3B are highlighted. A total of 19 MHR domain point mutations (Table S2) did not affect *Sm*.TRPM_PZQ_ responsiveness to PZQ. Truncation mutants at the COOH terminus are indicated by their location and functional effect on PZQ activation (red, inhibitory; green, no effect). (**B**) Homology model of *Sm*.TRPM_PZQ_ including docking pose of PZQ into the predicted binding site within the voltage-sensing ligand domain (VSLD, green) Pore domain (yellow) and TRP box domain (blue). (**C**) Hydrophobic cavity around cyclohexyl ring of PZQ. (**D**) PZQ and its non-covalent hydrogen bonding interactions (magenta) with binding site residues. (**E**) Hydrophobic cavity around aromatic ring of PZQ including ligand-sidechain and intra-residue hydrogen bond interactions (magenta). Single-letter code for amino acid residues: A, Ala; C, Cys; D, Asp; E, Glu; F, Phe; G, Gly; H, His; I, Ile; K, Lys; L, Leu; M, Met; N, Asn; P, Pro; Q, Gln; R, Arg; S, Ser; T, Thr; V, Val; W, Trp; and Y, Tyr.

Structures of human TRPM8 (*h*TRPM8) reveal that the transmembrane region (voltage sensor-like domain (VSLD), transmembrane helices S1-S4), encompasses a conformationally malleable binding pocket capable of accommodating diverse chemotypes (*15, 21*). This is relevant as (*S*)-PZQ has been shown to activate human TRPM8 (*h*TRPM8), albeit at a high micromolar range (EC_50_ ~20μM (*22*)). Using *h*TRPM8 structures as a template, point mutants of 11 known *h*TRMP8 pocket-lining residues were generated. These mutants impaired *h*TRPM8 activation in response to various ligands (Fig. S4). Overall, the pattern of *h*TRPM8 responsiveness to (*S*)-PZQ across this panel of *h*TRPM8 mutants most closely resembled responses with the menthol-related cooling agent WS-12. These data suggest (*S*)-PZQ activates *h*TRPM8 by interaction with the canonical *h*TRPM8 transmembrane ligand binding pocket. Does an equivalent PZQ binding pocket exist in the transmembrane region of *Sm*.TRPM_PZQ_? Five residues (Arg^1681^, Asn^1388^, Tyr^1392^, Arg^1514^ Tyr^1678^) identical to residues lining the *h*TRPM8 binding pocket were mutated within *Sm*.TRPM_PZQ_ (Fig. 2A). Four of these five mutants inhibited PZQ-evoked activation of *Sm*.TRPM_PZQ_ (Fig. 2A and S3B). In contrast, neighboring point mutations within the same regions did not prevent PZQ-evoked *Sm*.TRPM_PZQ_ activation (Table S4). These data suggest the existence of a TRPM8-like ligand binding site within the transmembrane region of *Sm*.TRPM_PZQ_.

Molecular modelling was employed to characterize the PZQ binding site. A homology model of the entire transmembrane region of *Sm*.TRPM_PZQ_ was built based from three known TRPM structures (Fig. 2B). Independent prediction tools identified a common binding site within the VSLD (Movie S1), in close proximity to the known *h*TRPM8 transmembrane pocket (Fig. S5). Docking simulations with (*R*)-PZQ revealed PZQ bridged two hydrophobic cavities separated by a region of polar contacts. The cyclohexyl ring of PZQ projected into one hydrophobic cleft defined at its inner apex by three hydrophobic residues – Phe^1395^ (S1), Trp^1421^ (S2) and Leu^1425^ (S2) – with residues Tyr^1392^ (S1) and Pro^1686^ (TRP) situated more peripherally around this ring (Fig. 2C). Within the polar region, the cyclohexyl carbonyl of PZQ forms a hydrogen bond with Arg^1514^ (S4-S5 linker) and the scaffold carbonyl of PZQ interacts with Asn^1388^, Tyr^1678^ and Arg^1681^ via hydrogen bonding (Fig. 2D). The second hydrophobic cavity accommodates the aromatic ring system of PZQ and is framed by six residues, Trp^1451^, Asn^1452^, Leu^1454^, Asp^1455^, Arg^1514^ and Tyr^1517^ (Fig. 2E). A salt bridge between Asp^1455^ and Arg^1514^ shaped this cavity, in which the unsaturated part of the PZQ scaffold interacted with Tyr^1517^ (S4-S5 linker) via π-π stacking and with Arg^1514^ (S4-S5 linker) via cation-π interactions.

The impact of alanine mutagenesis of all 23 pocket-lining residues lying within a 5Å radius of the binding pose was examined (Fig. 3A and Table S3). Mutation of 20 of these 23 residues decreased PZQ sensitivity (Fig. S6A and S6B), with many point mutants displaying a complete loss of PZQ-evoked activity (Table S3). In contrast, alanine mutagenesis of an additional 26 amino acid residues localized away from the predicted binding pocket, minimally impacted PZQ activation of *Sm*.TRPM_PZQ_ (Fig. S6C and Table S4). The localization of all 49 residues relative to the binding pocket is summarized in Figure S6.

**Fig. 3.**
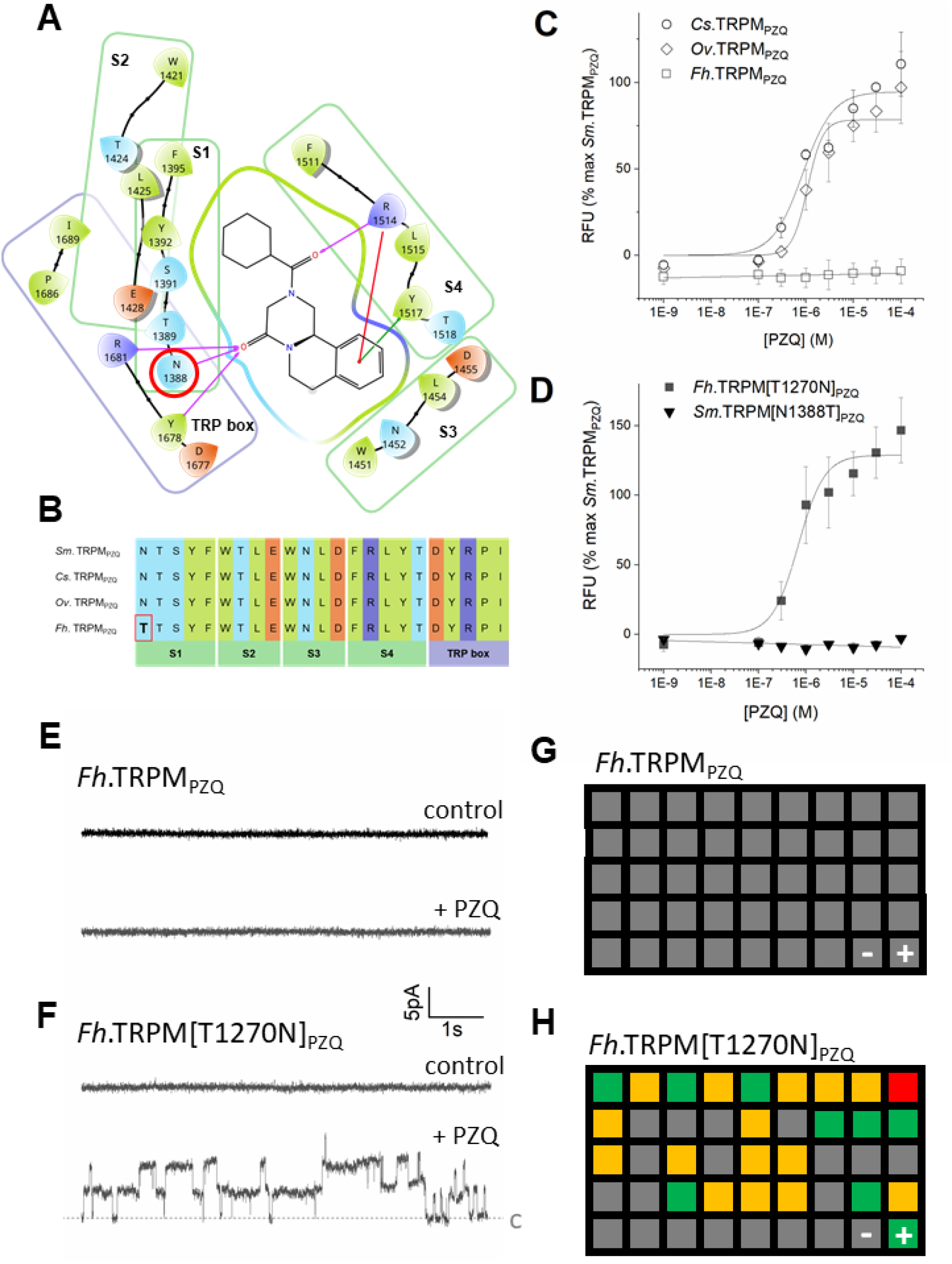
Analysis of PZQ binding pocket in trematode TRPM_PZQ_ channels. (**A**) Ligand-Receptor interaction map of (*R*)-PZQ with *Sm*.TRPM_PZQ_ showing amino acid residues within a 5Å radius and colored by properties (yellow: hydrophobic, blue: polar, purple: positively charged, orange: negatively charged). Interactions are highlighted by arrows (purple: hydrogen bonding, red: cation-π, green: π-π stacking). (**B**) Alignment of 23 binding pocket residues identified in *Sm*.TRPM_PZQ_ with other trematode TRPM_PZQ_ channels (*Clonorchis sinensis, Cs*.TRPM_PZQ_; *Opisthorchis viverrini, Ov*.TRPM_PZQ_ and *Fasciola hepatica, Fh*.TRPM_PZQ_). Sequence data is from genomic annotations (*Cs*.TRPM_PZQ_, GAA53128.1; *Ov*.TRPM_PZQ_, OON13534.1*; Fh*.TRPM_PZQ_, THD26109.1). (**C**) Concentration response relationship for TRPM_PZQ_ from different liver flukes. (**D**) Concentration response curves for reciprocal binding pocket mutants of *Schistosoma mansoni* and *Fasciola* TRPM_PZQ_. (**E & F**) Current traces recorded at +60 mV in HEK293 cells co-transfected with (E) *Fh*.TRPM_PZQ_ and GFP, or (F) *Fh*.TRPM_PZQ_(Thr^1270^→Asn) and GFP, in the cell-attached configuration after addition of 0.1% DMSO (top, control) and 10 μM PZQ (bottom). c, closed state. (**G** & **H**) Graphical heatmap of the effects of each of the 43 analogues on (G) *Fh*.TRPM_PZQ_ and (H) *Fh*.TRPM[T1270N]_PZQ_ activation. Each analogue is numbered left to right (1 through 43) in the heatmap and indexed to Table S6, with responses to vehicle (-) and PZQ (+) indicated (bottom right). Grid is color-coded as Figure 1C, with colors describing analogs with EC_50_s <1μM (green), 1-10μM (yellow), >10μM (red) or inactive (grey).

This mutational analysis verified key aspects of the binding model. First, mutagenesis of channel residues predicted to interact with the carbonyl groups of PZQ (Fig. 2D and 3A) either caused a complete loss (Asn^1388^→Ala, Arg^1514^→Ala, Tyr^1678^→Ala) or significant reduction (Arg^1681^→Ala) in *Sm*.TRPM_PZQ_ sensitivity (Table S3). Second, mutagenesis of residues in the hydrophobic binding cavity surrounding the cyclohexyl ring of PZQ led to complete loss of PZQ-evoked activity (Fig. 2C and Table S3). This hydrophobic cavity is tightly framed by residues from the S1 (Tyr1^392^, Phe^1395^) and S2 helices (Trp^1421^, Leu^1425^) as well as the TRP domain (Pro^1686^, Fig. 2C). Mutation of Leu^1425^, predicted to lie above the cyclohexyl ring of PZQ (Fig. 2C) to either bulkier (Leu^1425^→Phe) or smaller residues (Leu^1425^→Ala) decreased PZQ sensitivity (Table S3). This is consistent with the analogue SAR analysis defining the cyclohexyl ring of PZQ as a key part of the pharmacophore and poorly tolerable to substitution. This requirement for activity conflicts with the incorporation of a polar hydroxyl group onto the cyclohexyl ring during phase 1 metabolism of PZQ (Fig. 1A and S2E). The major trans-(*R*)-4-OH PZQ metabolite (analog **21**, Fig. 1A), exhibits a ~36-fold reduction in activity at *Sm*.TRPM_PZQ_ (Tables S1 & S5). Efforts to design more potent and metabolically-stable analogs to allow for lower dosing and sustained exposure may prove difficult in light of these observed structural requirements. Third, mutagenesis of residues framing the second hydrophobic cavity around the aromatic ring of PZQ led to complete loss (Trp^1451^, Arg^1514^, Tyr^1517^) or decreased PZQ-evoked activity (Leu^1454^, Asn^1452^; Table S3). Mutation of Asp^1455^→Ala, which electrostatically interacts with Arg^1514^ to define this pocket (Fig. 2E) also ablated PZQ sensitivity (Table S3). Interactions that orientate the (*R*)-PZQ cyclohexyl ring within the first hydrophobic pocket serve to preserve optimal π-π stacking of the aromatic ring with Tyr^1517^ (S4) in this second hydrophobic pocket. (*R*)-PZQ therefore bridges two hydrophobic cavities, occupied by opposite ends of the PZQ molecule, to cause potent channel activation. For (*S*)-PZQ, the binding orientation is not optimal (Fig. S2D) providing a structural basis for the ~50-fold difference in EC_50_s of the enantiomers at *Sm*.TRPM_PZQ_. Even subtle changes in structure as seen with the veterinary drug epsiprantel (Fig. 1A, analog **35**), which contains a seven-membered azepine ring, alter ligand conformation (Fig. S2F) and result in lower potency. Therefore, both modeling and mutagenesis data converge to establish that (*R*)-PZQ engages *Sm*.TRPM_PZQ_ via a transmembrane binding pocket to cause channel activity, Ca^2+^ influx and worm contraction.

PZQ is used to treat other parasitic flatworm infections, such as clonorchiasis and opisthorchiasis (*23*). The sequences of the TRPM homologs in the liver flukes *Clonorchis sinensis* and *Opisthorchis viverrini* displayed complete conservation of the 23 residues that most closely circumscribe the PZQ binding site (Fig. 3B). Heterologous expression of *Cs*.TRPM_PZQ_ and *Ov*.TRPM_PZQ_ conferred PZQ-evoked Ca^2+^ signals in transfected HEK293 cells. The observed sensitivities of the fluke TRPM_PZQ_ channels to PZQ (*Cs*.TRPM_PZQ_, EC_50_=0.85±0.22μM; *Ov*.TRPM_PZQ_, EC_50_=1.07±0.33μM; Fig. 3C) were similar to *Sm*.TRPM_PZQ_. One parasitic flatworm infection that is refractory to PZQ treatment is fascioliasis, a disease of considerable clinical and veterinary importance worldwide. The *Fasciola hepatica* TRPM_PZQ_ channel (*Fh*.TRPM_PZQ_) did not respond to PZQ (Fig. 3C), unlike the TRPM_PZQ_ channels in these other PZQ-sensitive flukes. Sequence analysis of *Fh*.TRPM_PZQ_ revealed a single amino acid difference in the predicted PZQ binding pocket compared with PZQ-sensitive TRPM_PZQ_ channels where a threonine residue in S1 of *Fh*.TRPM_PZQ_ replaced an asparagine residue present in the other homologs (Fig. 3B). This difference reflects a single nucleotide difference in the genomic sequence of these flatworms (‘ACT’ encoding threonine in *Fasciola*, ‘AAT’ encoding asparagine in *Schistosoma*). The resulting asparagine residue in the PZQ-sensitive channels is predicted to be a critical residue that interacts with the core carbonyl of PZQ (Fig. 2D and 3A).

To test the impact of substitution at this residue, reciprocal point mutants were generated in *Sm*.TRPM_PZQ_ (Asn^1388^→Thr) and *Fh*.TRPM_PZQ_(Thr^1270^→Asn). The asparagine to threonine mutation in the schistosome channel caused a loss of PZQ-evoked activity (Fig. 3D). Reciprocally, the threonine to asparagine mutation in the *Fasciola* channel (Thr^1270^→Asn) caused a gain of responsiveness to PZQ (Fig. 3D), with the mutant *Fasciola* channel being robustly activated by PZQ (EC_50_=0.55±0.1μM). The properties of the *Fasciola* channels were then assessed electrophysiologically. Whereas PZQ failed to activate currents in cells expressing wild type *Fh*.TRPM_PZQ_ (Figure 3E), addition of PZQ activated step-like changes in transmembrane currents in cells expressing *Fh*.TRPM_PZQ_(Thr^1270^→Asn), consistent with PZQ-evoked single channel activity (Figure 3F). Therefore, two orthogonal assays (Ca^2+^ imaging and electrophysiology) demonstrate a single point mutation in the *Fasciola* binding pocket results in a gain of sensitivity to PZQ. These data reveal a molecular basis for the natural insensitivity of *Fasciola* to PZQ chemotherapy.

Finally, we screened the 43 analogues used to define structure-activity relationships at *Sm*.TRPM_PZQ_ at *Fh*.TRPM_PZQ_. No activity was resolved in response to any analogue at the wild type channel (Fig. 3G). In contrast, in cells expressing *Fh*.TRPM_PZQ_(Thr^1270^→Asn), several analogues displayed activity (Fig. 3H). The structure-activity pattern was identical at both schistosome and liver fluke channels (compare Fig. 1C with Fig 3H), with the same 24 analogs that activated *Sm*.TRPM_PZQ_ also activating *Fh*.TRPM_PZQ_(Thr^1270^→Asn). *Fh*.TRPM_PZQ_(Thr^1270^→Asn) displayed ~3-fold higher sensitivity to the active analogs, consistent with a higher sensitivity of *Fh*.TRPM_PZQ_(Thr^1270^→Asn) to PZQ compared with *Sm*.TRPM_PZQ_.

In conclusion, this study identifies a binding site for (*R*)-PZQ in a juxta-membrane cavity within the VSLD domain of TRPM_PZQ_, a broadly conserved parasitic flatworm ion channel. Identification of a *h*TRPM8-like binding pocket within an invertebrate TRPM2-like channel architecture was surprising, but highlighting how little we know about the properties of flatworm TRPs (*24*) which have specialized along evolutionary trajectories distinct from human TRP channels. Mutagenesis of the PZQ binding site in *Sm*.TRPM_PZQ_ demonstrates that single point mutations ablate responsiveness to PZQ, and the properties of wild-type *Fh*.TRPM_PZQ_ evidence a naturally occurring TRPM_PZQ_ channel that is insensitive to PZQ. The significance of these changes in context of reports of decreased clinical effectiveness of PZQ in the field (*25, 26*), or the lower sensitivity of isolated schistosome strains to PZQ (*27, 28*), is important to monitor given the critical need to preserve the effectiveness of PZQ for treating this burdensome neglected tropical disease. Definition of the (*R*)-PZQ binding site in this TRP channel now provides impetus to design novel therapies that exploit the druggability of TRPM_PZQ_ in various disease-causing parasites.

## Non-Author Contributions

DM acknowledges Dr. Andreas Waechtler, Jeremy Maurin, Mathilde Dreyer, Philipp Rietsch, Marie Emering, Fabian Gramain and Minas Stefanou for their contributions to chemical synthesis.

## Author Contributions

Experiments were performed by SKP, NY, CR. Chemical synthesis of PZQ analogues was directed by DM. Computational chemistry and molecular modeling were performed by LF and FR. Structure-activity relationship studies were directed by TS. SKP, LF, TS and JSM designed experimental analyses, TS and JSM wrote the paper with help from all authors.

