## Supplementary Material for "Mechanism of praziquantel action at a parasitic flatworm ion channel"

**This PDF file includes**

Materials and Methods
Figs. S1 to S6
Tables S1 to S6
Caption for Movie S1

**Other Supplementary Materials for this manuscript include the following:**

Movie S1

**Materials and Methods**.

**Reagents.** Cell culture reagents were from Invitrogen. Chemical reagents were from Sigma or Thermofisher. PZQ derivatives were synthesized by published procedures (referenced in Table S1), or by the protocols detailed below. The human TRPM8 clone was from R&D Systems (RDC0188).

**Cell Culture and transfection.** HEK293 cells were purchased from ATCC (CRL-1573.3) and verified by STR profiling. All cultures were determined negative for mycoplasma contamination. Cells were used for heterologous expression of either *Sm*.$TRPM_{PZQ}$ or specified mutants at low passage numbers (<20). HEK293 cells were cultured in DMEM supplemented with 10% fetal bovine serum (FBS), penicillin (100 units/ml), streptomycin (100μg/ml), and L-glutamine (290μg/ml). For transfection, plasmids were transiently transfected into HEK293 cells using Lipofectamine 2000 at a density of $3x10^6$ cells per dish (100mm), and transfected cells replated in multiwell plates for $Ca^{2+}$ imaging assays. Point mutants prepared in the wild type *Sm*.$TRPM_{PZQ}$ backbone were confirmed by sequencing.

**Adult schistosome mobility assays***.* Adult schistosomes were recovered after dissection of the mesenteric vasculature in female Swiss Webster mice previously infected (~49 days) with *Schistosoma mansoni* cercariae (NMRI strain) by the Schistosomiasis Resource Center at the Biomedical Research Institute (Rockville, MD). Animal husbandry and experimentation followed ethical regulations approved by the MCW IACUC committee. After isolation, harvested schistosomes were washed in RPMI 1640 supplemented with HEPES (25mM), 5% heat inactivated FBS (Gibco) and penicillin-streptomycin (100 units/mL), separated into 6-well plates (3 males and 3 females per well) and incubated overnight (37°C/5% $CO_2$). The following day, worm movement assays in response to drug exposures were performed (~3 male and 3 female, ~3ml of media per well in a six well dish). Recordings of worm motility before and after addition of various PZQ analogues (500nM, 5μM and 50μM), were captured using a Zeiss Discovery v20 stereomicroscope coupled to a QiCAM 12-bit cooled color CCD camera controlled by Metamorph imaging software. Recordings were made at 5 mins, 1 day and 2 days after continual drug exposure. Data were recorded as a sequential image stack (60s duration, 100ms exposure every 2s) and analyzed using the following workflow. A maximal intensity projection of the entire stack was made, which was subsequently filtered by a single fast Fourier transform (FFT) blur algorithm to provide a single image representation of worm movement over the entire duration of the video. This image was thresholded to identify the total pixel area covered by individual worm movements during the recording time, and the percentage of the thresholded area within the fixed imaging frame (1392x1040 pixels) resolved. This area represented the proportion of pixels occupied by movement of an individual worm throughout the entire recording (typically ~6% of the field of view under control conditions at room temperature). For clarity of data display, an edge detected filer was applied to the maximal intensity projection, colorized and merged with a single frame video image to show a single snapshot of the worms overlaid with their resulting movement arcs over the entire recording period. This output was averaged for each experiment and then between assays, normalized to values resolved with control and PZQ-exposed worms. This value was reported as mean±sem for ≥3 independent experiments, defined as separate isolations of worms from different cycles of infection.

**$Ca^{2+}$ imaging assays.** $Ca^{2+}$ imaging assays for quantifying the activity of PZQ analogues at *Sm*.$TRPM_{PZQ}$ as well as profiling other $TRPM_{PZQ}$ channels and *h*TRPM8 were performed using a Fluorescence Imaging Plate Reader ($FLIPR^{TETRA}$, Molecular Devices). Briefly, HEK293 cells (naïve or transfected) were seeded in black-walled clear-bottomed poly-D-lysine coated 96-well (50,000 cells/well) or 384-well plates (20,000 cells/well, Corning) in DMEM growth media supplemented with 10% FBS. After 24hrs, growth medium was removed, and cells were loaded with a fluorescent $Ca^{2+}$ indicator (Fluo-4 NW dye, Invitrogen) by incubation (100μl per well, 30mins at 37°C, followed by an additional 30mins at room temperature) in Hanks' balanced salt solution (HBSS) assay buffer containing probenecid (2.5 mM) and HEPES (20mM). Drug dilutions were prepared in assay buffer, without probenecid and dye. After indicator loading, the fluorometric $Ca^{2+}$ assay was performed at room temperature. Basal fluorescence was monitored for 20s, then 25µl (96-well plate) of each drug was added, and the signal (raw fluorescence units) monitored for an additional 250s. For quantitative analyses, peak fluorescence in each well was normalized to maximum-fold increase over baseline. Changes in fluorescence amplitude were analyzed using the sigmoidal dose-response function in Origin. Results are collated from a minimum of three biological replicates (independent transfections) for each construct. Each individual assay comprised three technical replicates on a single multi-well plate.

**Electrophysiology.** As an orthogonal assay to assess $TRPM_{PZQ}$ properties, electrophysiological analyses were performed in $Ca^{2+}$ free solutions. Single channel current recordings were made in cell-attached mode from HEK293 cells plated onto glass coverslips. Cells were co-transfected with plasmids encoding GFP and *Fh.*$TRPM_{PZQ}$ or *Fh.*$TRPM_{PZQ}$[T1270N]. After 24 hours coverslips were secured into a recording chamber on a Olympus BX51WI upright microscope. The bath solution contained (in mM) 145 NaCl, 10 HEPES, 1 EGTA, pH 7.4-NaOH (310-315 mOsm with sucrose). Pipette solution was (in mM) 140 LiCl, 10 HEPES, 1 EGTA, pH 7.4-LiOH (280-285 mOsm). Patch pipettes were made from borosilicate glass (BF150-110-10, Sutter Instrument, Novato, CA) pulled on a vertical puller (Narishige, Amityville, NY, Model PC-10, resistances 8–10 MΩ). Recordings were performed using a MultiClamp 700B amplifier and Digidata 1440A digitizer (Molecular Devices, Sunnyvale, CA), filtered with an 8-pole Bessel low pass filter at 1 kHz, and analyzed through Clampfit 10 software. All recordings were made at the room temperature. PZQ and vehicle (DMSO) solutions were added directly to the recording chamber.

**SAR activity models, homology modelling and molecular docking.** SAR analyses were performed using Cresset's Forge (version 10.6.0) (*29*). An activity model was built using the Activity Atlas module to investigate interaction regions. Calculations were performed with the Schrödinger Suite molecular modelling package (release version 2019-04) utilizing program default parameters unless otherwise reported. The homology model was built with Prime (release version 2019-04) using a shortened section (residues 1100 to 1800) of the *Sm*.$TRPM_{PZQ}$ sequence (*12*). Three Protein Data Bank (PDB) structures were used to build a consensus homology model: 6CO7 (TRPM2), 6BCJ (TRPM4) and 6NR3 (TRPM8). Template structures were prepared by the default preparation protocol of the Protein Preparation Wizard, including hydrogen-bond optimization at pH7 and a protein structure minimization with the force field OPLS3e (*30*). Potential binding sites were predicted with SiteMap (*31*) and the web-based tool DogSiteScorer (*32*). (*R*)-PZQ was prepared with LigPrep and docked into the selected binding pocket using the Induced Fit Docking protocol. Interactions of the top-ranked docking pose were further investigated.

***In vitro* metabolism assays in liver microsomes**. The metabolic stability assay was performed in duplicate in a 96-well microtiter plate. The test compounds (0.1µM) were incubated (37 °C) in mouse and pooled human liver microsomes (final protein concentration of 0.4 mg/mL; XenoTech, Lenexa, KS) suspended in 0.1 M phosphate buffer (pH 7.4) for predetermined time points, in the presence and absence of the cofactor NADPH (1 mM). The reactions were quenched by the addition of ice-cold acetonitrile containing internal standard (carbamazepine, 0.0236µg/mL). The samples were centrifuged and the supernatant was filtered and analyzed by means of LC-MS/MS (Agilent Rapid Resolution HPLC, AB SCIEX 4500 MS). The relative loss of parent compound over time was monitored and plots were prepared for each compound of concentration versus time to determine the first order rate constant for compound depletion.

**Chemical Synthesis**. All solvents and reagents were of pharmaceutical grade, acquired from commercial suppliers or internally from the Merck production and used without additional purifications. Thin-layer chromatography was conducted on E. Merck silica gel 60 F-254 precoated aluminium plates; detection was performed using a UV short-wave lamp. Column chromatography was carried out at atmospheric pressure using Silica Gel 60 (70-230 mesh, from E. Merck). HPLC analysis was performed on VWR EliteLaChrom units and different chiral columns, depending on each compound. For Praziquanamine derivatives a Chiralpak AS-H column was used. For Praziquantel derivatives a Chiralpak IA column was used. The NMR spectra were recorded on Brucker Advance II spectrometers operating at 300, 400 or 500 MHz for $^{1}H$ NMR and at 100 MHz for $^{13}C$ NMR. Mass spectrometry was performed on an Agilent Triple Quad spectrometer. Melting points were determined on a DSC 3+ Mettler Toledo unit.

**Synthesis of (*R*)-Praziquantel homologues:**

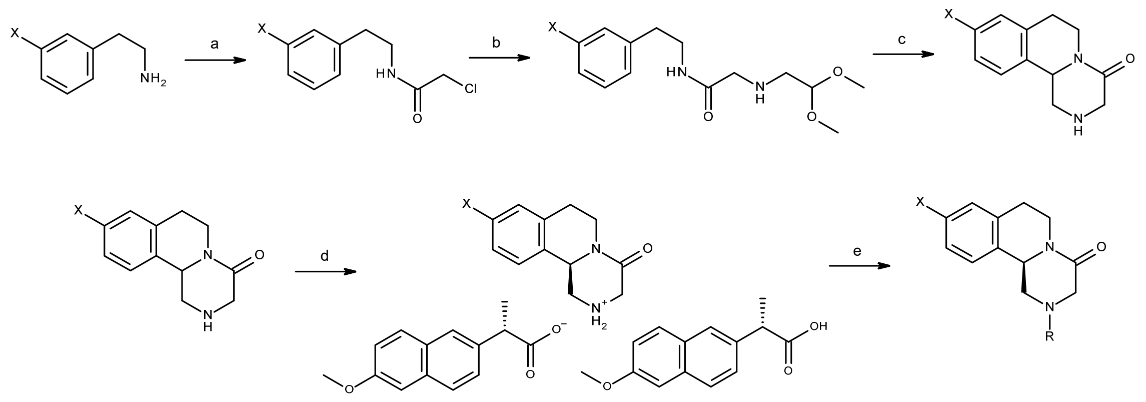


a. $ClCH_2COCl$, NaHCO3, DCM; b. $NH_2CH_2CH(OCH_3)_2$, NaOH, $H_2O$; c. $H_2SO_4$, DCM; d. Naproxen, iPrOH, $H_2O$; e. 1. MeSO3H, $H_2O$, 2. RCOCl, NaOH, DCM

## X=H :

Synthesis of racemic Praziquanamine already described.
*Kim et al., Heterocycles 1998, 42, 2279; Tetrahedron 1998, 54, 7395; DE3324532; US4497952*

Resolution of racemic Praziquanamine with Naproxen already described.
*D. Maillard, A. Waechtler, J. Maurin, E. Wakaresko, C. Jasper, WO2016078765*

### (*R*)-1,2,3,6,7,11b-Hexahydro-pyrazino[2,1-a]isoquinolin-4-one-di-(*S*)-(6-methoxy-naphtalen-2-yl)-propionic acid salt

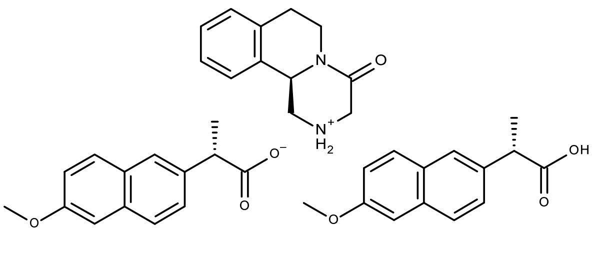


520g (2.54mol) racemic (*R*)-Praziquanamine and 585g (2.54mol) Naproxen are heated under stirring in a solvent mixture made from 2-propanol (4010g) and water (1005g) up to 65°C until complete dissolution. The resulting solution is stirred and cooled down to -19°C. When the target temperature is met, the precipitate is filtered off, the filter cake is washed two times with 780g of cold (-20°C) 2-propanol. After drying (24h at 50°C under vacuum) 746.7g (1.13mol) of (*R*)-Praziquanamine/Naproxen salt (R/SS) are obtained as a white solid (44.4% yield) with an enantiomeric excess of 98.6%(R).
$C_{40}H_{42}N_2O_7$ (662.77g.mol$^{-1}$), m.p. 157-159°C. $^1$H NMR($CD_3OD$) : 6.14-6.19 (m, 6H$_{arom}$), 5.87 (dt, J=8, 2Hz, 2H$_{arom}$), 5.62-5.72 (m, 6H$_{arom}$), 5.56 (ddd, J=8, 4, 2Hz, 2H$_{arom}$), 3.31 (dd, J=8 ,4 Hz, 1H), 3.14-3.18 (m, 1H), 2.35 (s, 6H), 2.28 (dd, J=12, 8Hz, 2H), 2.23 (dd, J=8 ,4Hz, 1H), 1.98 (AB, J=18Hz, 2H), 1.77-1.79 (m, 4H), 1.21-1.38 (m, 4H), 0.01 (d, J=8 Hz, 6H). $^{13}$C NMR($CD_3OD$): 177.2 (s, 2CO$_2$H), 167.5 (s, 1C=O), 157.6 (s, 2$\underline{C}$OMe), 136.2 (s, 2qC$_{arom}$), 134.5 (s, 1qC$_{arom}$), 133.7 (s, 2qC$_{arom}$), 133.6 (s, 1qC$_{arom}$), 129.0 (s, 2qC$_{arom}$), 128.8 (s, 1C$_{arom}$), 128.7 (s, 2C$_{arom}$), 126.7 (s, 1C$_{arom}$), 126.6 (s, 2C$_{arom}$), 126.4 (s, 1C$_{arom}$), 125.8 (s, 2C$_{arom}$), 125.4 (s, 2C$_{arom}$), 124.5 (s, 1C$_{arom}$), 118.4 (s, 2C$_{arom}$), 105.2 (s, 2C$_{arom}$), 55.7 (s, 1CH), 54.3 (s, 2OCH$_3$), 48.3 (s, 1CH$_2$), 48.1 (s, 1CH$_2$), 45.3 (s, 2CH), 38.8 (s, 1CH$_2$), 28.2 (s, 1CH$_2$), 17.6 (s, 2CH$_3$).

### (*R*)-1,2,3,6,7,11b-Hexahydro-pyrazino[2,1-a]isoquinolin-4-one

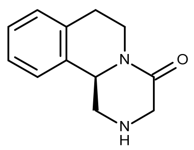


60g (*R*)-Praziquanamine/Naproxen salt (0.09mol) are suspended in water (180g). At 25°C and under stirring, MeSO3H (6.47 mL, 0.10mol) is slowly added. After stirring for 2h, the suspension is filtered and the cake washed with water (100g). 32% NaOH (13g) is added to the combined mother and wash liquors and the resulting aqueous mixture is extracted with DCM (4x60mL). The solvent of the combined organic phases is removed under reduced pressure to afford 17.72g (0.088mol) of (*R*)-Praziquanamine as a white-off solid (96.8% yield) with an enantiomeric excess of 98.5%.

$C_{12}H_{14}N_2O$ (202.25g.mol$^{-1}$), m.p. 118.5°C. $^1$H NMR(DMSO-d$_6$): 7.23-7.34 (m, 4H$_{arom}$), 6.05 (bs, NH), 5.05 (dd, J=12, 4Hz, 1H), 4.60 (dt, J=8, 4Hz, 1H), 4.13 (dd, J=12, 2Hz, 1H), 3.76 (m, 2H), 3.14 (m, 1H), 2.93 (m, 1H), 3.14 (m, 2H). $^{13}$C NMR(DMSO-d$_6$): 168.1 (s, 1C=O), 134.6 (s, 1qC$_{arom}$) 133.9 (s, 1qC$_{arom}$), 129.8 (s, 1C$_{arom}$), 126.6 (s, 1C$_{arom}$), 126.3 (s, 1C$_{arom}$), 124.5 (s, 1C$_{arom}$), 56.0 (s, 1CH), 48.6 (s, 1CH$_2$), 48.5 (s, 1CH$_2$), 38.8 (s, 1CH$_2$), 28.3 (s, 1CH$_2$). MS (EI): m/z (%): 202 (40) [M]$^+$, 173 (67), 145 (100), 131 (58), 117 (22), 103 (10), 77 (9), 43 (10).

**(*R*)-4-Oxo-1,3,4,6,7,11b-hexahydro-pyrazino[2,1-a]isoquinoline-2-carboxylic acid *tert*-butyl ester (23)**

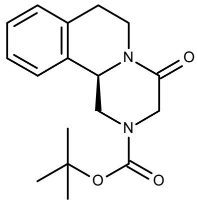


28.06g Di-*tert*-butyl-dicarbonate (0.128mol) are dissolved in DCM (40mL) and stirred at room temperature under nitrogen. A solution of 13.0g (*R*)-Praziquanamine (0.064mol) dissolved in DCM (40mL) was added dropwise. After 4 hours the reaction mixture is washed with an aqueous solution of HCl (2.5 %-w, 2x100mL) and with water (100mL). The organic phase is dried over $Na_2SO_4$ for 1h, filtered and the solvent evaporated. The crude product is purified by column chromatography (EtOAc/*n*-Hept 2/3 and 2/1) to afford the target compound (16.54g, 0.055mol, 85.1% yield) as a white solid.

$C_{17}H_{22}N_2O_3$ (302.37g.mol$^{-1}$), m.p. 118-121°C. $^1$H NMR (DMSO-d$_6$): 7.16-7.28 (m, 4H$_{arom}$), 4.81-4.88 (m, 2H), 4.63-4.76 (m, 1H), 4.50 (AB, *J*= 18Hz, 1H), 3.87 (AB, *J*= 18Hz, 1H), 2.74-3.03 (m, 4H), 1.51 (s, 9H). $^{13}$C NMR (DMSO-d$_6$): 165.3 (s, 1C=O), 153.8 (s, 1C=O), 135.0 (s, 1qC$_{arom}$), 132.8 (s, 1qC$_{arom}$), 129.3 (s, 1C$_{arom}$), 127.4 (s, 1C$_{arom}$), 126.8 (s, 1C$_{arom}$), 125.3 (s, 1C$_{arom}$), 56.0 (s, 1qC), 55.3 (s, 1CH), 47.8 (s, 1CH$_2$), 46.5 (s, 1CH$_2$), 38.9 (s, 1CH$_2$), 28.8 (s, 1CH$_2$), 28.4 (s, 3CH$_3$). MS (EI): m/z (%): 302 (1) [M]$^+$, 246 (28), 201 (31), 173 (19), 145 (20), 132 (37), 57 (100).

**X=F:**

**2-Chloro-N-[2-(3-fluoro-phenyl)-ethyl]-acetamide**

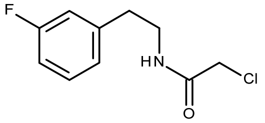

5g 2-(3-Fluoro-phenyl)-ethylamine (34.85mmol) and 4.39g sodium bicarbonate (52.27mmol) are dissolved in DCM (100mL). After cooling down the stirred solution at a temperature between -5°C and 0°C, a solution of 3.36mL chloro-acetyl chloride (41.82mmol) in DCM (30mL) is slowly added dropwise. The reaction mixture is then stirred for 1.5h within a temperature range from -5°C to 5°C before addition of water (50mL). After settling for 15 min, the phases are separated, the organic phase is washed twice with a 10% aqueous solution of sodium bicarbonate (100mL) and finally with water until the pH is in a range of 5-7. The solvent is removed under vacuum affording the target compound as a white solid (7.50g, 34.77mmol, 99.8% yield).

$C_{10}H_{11}ClFNO$ (215.65g.mol$^{-1}$), m.p. 70-75°C. $^{1}$H NMR ($CDCl_3$): 7.27-7.34 (m, 1$H_{arom}$), 6.91-7.03 (m, 3$H_{arom}$), 6.62 (bs, 1NH), 4.05 (s, 2H), 3.59 (dd, *J*=12, 6Hz, 2H), 2.88 (t, *J*=6Hz, 2H). $^{13}$C NMR ($CDCl_3$): 165.8 (s, 1C=O), 153.8 (d, *J*=245Hz, 1q$C_{arom}$), 140.8 (d, *J*=8Hz, 1q$C_{arom}$), 130.2 (d, *J*=8Hz, 1$C_{arom}$), 124.3 (d, *J*=3Hz, 1$C_{arom}$), 115.6 (d, *J*=21Hz, 1$C_{arom}$), 113.7 (d, *J*=21Hz, 1$C_{arom}$), 42.6 (s, 1$CH_2$), 40.7 (s, 1$CH_2$), 35.2 (d, *J*=3Hz, 1$CH_2$). MS (EI): m/z (%): 215 (7) [M]$^+$, 180 (4), 122 (100), 106 (20), 96 (4), 83 (12), 77 (10), 57 (4), 49 (5), 30 (49).

**2-(2,2-Dimethoxy-ethylamino)-*N*-[2-(3-fluoro-phenyl)-ethyl]-acetamide**

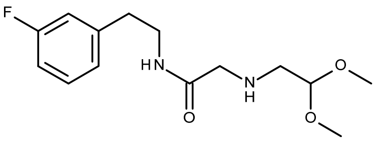

3.83g sodium Hydroxide (30.60mmol) are dissolved in water (100mL). After addition of 15.16mL 2,2-Dimethoxy-ethylamine (139.11mmol) at RT, the solution is heated up at 50°C under nitrogen and 6g 2-Chloro-N-[2-(3-fluoro-phenyl)-ethyl]-acetamide (27.82 mmol) are added portion wise. After dissolution of all solids, the mixture is kept at 50°C during 21h before removal of water and the excess of 2,2-Dimethoxy-ethylamine under vacuum. DCM (130mL) and water (100mL) are added to the residue which is dissolved under stirring at 25°C. The phases are subsequently separated, the organic layer is washed twice with water (50mL). The solvent is finally removed under vacuum affording the target compound as a slightly yellow oil (6.83g, 24.02mmol, 86.3% yield).
$C_{14}H_{21}FN_2O_3$ (284.33g.mol$^{-1}$). $^{1}$H NMR ($CDCl_3$): 7.33 (bs, 1NH), 7.21-7.28 (m, 1$H_{arom}$), 6.87-6.98 (m, 3$H_{arom}$), 4.28-4.32 (m, 1H), 3.52-3.56 (m, 2H), 3.36-3.41 (m, 1H), 3.35 (s, 6H), 3.21-3.28 (m, 2H), 2.84 (t, *J*=8Hz, 1H), 2.73-2.81 (m, 1H), 2.65 (d, *J*=8Hz, 2H). $^{13}$C NMR ($CDCl_3$): 171.4 (s, 1C=O), 162.9 (d, *J*=244Hz, 1q$C_{arom}$), 141.6 (d, *J*=7Hz, 1q$C_{arom}$), 130.0 (d, *J*=8Hz, 1$C_{arom}$), 124.3 (d, *J*=3Hz, 1$C_{arom}$), 115.6 (d, *J*=20Hz, 1$C_{arom}$), 113.3 (d, *J*=20Hz, 1$C_{arom}$), 103.5 (s, 1CH), 54.0 (s, 2$CH_3$), 52.3 (s, 1$CH_2$), 51.1 (s, 1$CH_2$), 39.7 (s, 1$CH_2$), 35.5 (s, 1$CH_2$). MS (EI): m/z (%): 284 (1) [M]$^+$, 252 (19), 209 (12), 152 (16), 123 (8), 118 (34), 109 (5), 86 (92), 75 (100), 58 (19), 42 (16), 30 (5).

**9-Fluoro-1,2,3,6,7,11b-hexahydro-pyrazino[2,1-a]isoquinolin-4-one**

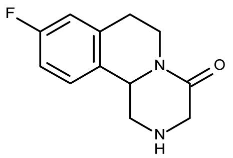


29.57g sulphuric acid (295.43 mmol) are dissolved in DCM (300mL) and stirred at 0°C under nitrogen. A solution of 21g 2-(2,2-Dimethoxy-ethylamino)-*N*-[2-(3-fluoro-phenyl)-ethyl]-acetamide (73.86mmol) in DCM (100mL) is then added dropwise keeping the temperature below 10-15°C. After complete addition, the temperature is increased to 25°C and the mixture is stirred for 6h before addition of water (400mL). After stirring for 30min, the phases are separated, the aqueous phase is washed with DCM (200mL). 27.69g of a 32% aqueous solution of sodium hydroxide (221.58 mmol) is added dropwise to the aqueous layer leading to a pH of 12-14. This mixture is extracted 3 times with DCM (100mL). The solvent of the combined organic phases is removed under vacuum. The residual orange oil is purified by column chromatography (pure AcOEt then AcOEt/MeOH 9/1) affording the target compound as red oil (10.11g, 45.90mmol, 62% yield).

$C_{12}H_{13}FN_2O$ (220.24g.mol$^{-1}$). $^{1}$H NMR (DMSO-d$_6$): 7.31 (dd, J=8, 6Hz, 1H$_{arom}$), 7.01-7.08 (m, 2H$_{arom}$), 4.71 (dd, *J*=12, 4Hz, 1H), 4.58-4.66 (m, 1H), 3.67 (dd, *J*= 12, 4Hz, 1H), 3.21-3.34 (m, 2H), 2.73-2.78 (m, 3H), 2.58 (dd, *J*=12, 10Hz, 1H). $^{13}$C NMR (DMSO-d$_6$): 167.4 (s, 1C=O), 161.0 (d, *J*=241Hz, 1qC$_{arom}$), 137.8 (d, *J*=7Hz, 1qC$_{arom}$), 131.7 (d, *J*=3Hz, 1qC$_{arom}$), 127.6 (d, J=8Hz, 1C$_{arom}$), 115.5 (d, *J*=21Hz, 1C$_{arom}$), 113.8 (d, *J*=21Hz, 1C$_{arom}$), 56.1 (s, 1CH), 50.1 (s, 1CH$_2$), 49.7 (s, 1CH$_2$), 38.1 (s, 1CH$_2$), 28.9 (s, 1CH$_2$). MS (EI): m/z (%): 220 (24) [M]$^+$, 191 (60), 163 (100), 149 (74), 135 (32), 122 (13), 109 (9), 101 (15), 96 (6), 75 (6), 43 (32).

**(11b*R*)-9-fluoro-1,2,3,6,7,11b-hexahydropyrazino[2,1-a]isoquinolin-2-ium-4-one;(2*S*)-2-(6-methoxy-2-naphthyl)propanoate;(2*S*)-2-(6-methoxy-2-naphthyl)propanoic acid**

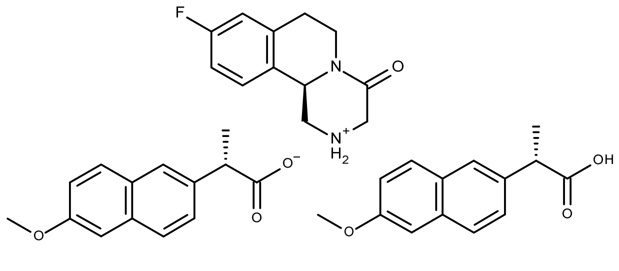


5.84g racemic fluorinated Praziquanamine (9-Fluoro-1,2,3,6,7,11b-hexahydro-pyrazino[2,1-a]isoquinolin-4-one (26.52mmol) and 6.11g Naproxen (26.52mmol) are dissolved in a mixture of Isopropanol (40mL) and water (20mL) at 65°C. After complete dissolution, the solution is cooled down to 0°C. The resulting suspension is kept under stirring at 0°C during 40 min before filtration of the orange precipitate. The filter cake is washed 3 times with cold Isopropanol (50 mL). After drying under vacuum at 40°C, the target salt is obtained as white solid (5.05g, 7.42mmol, 28% yield).
$C_{40}H_{41}FN_2O_7$ (680.76g.mol$^{-1}$).

**(*R*)-2-Cyclohexanecarbonyl-9-fluoro-1,2,3,6,7,11b-hexahydro-pyrazino[2,1-a]isoquinolin-4-one (30)**

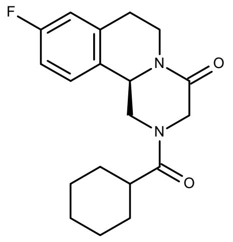

5.05g fluorinated (*R*)-Praziquanamine/Naproxen salt (7.40mmol) are suspended in water (60mL) under stirring. 0.53mL methanesulfonic acid (8.14mmol) are added dropwise over 10 min, the mixture is then further stirred for 2h before filtration and the wet cake is washed 3 times with water (60mL). To the combined aqueous mother and wash liquors, DCM (150mL) is added and the biphasic mixture is stirred at 25°C under nitrogen. A 32% aqueous solution of sodium hydroxide is added to reach a basic pH of 10. A solution of 1.05mL cyclohexanecarboxlic acid chloride (7.77mmol) in DCM (50mL) and 0.97g of a 32% aqueous solution of sodium hydroxide (7.77mmol) are subsequently added dropwise simultaneously keeping the pH between 9 and 11. After complete addition, the reaction mixture is stirred for 2h at RT and finally allowed to settle. The phases are separated, the aqueous one is extracted 3 times with DCM (50mL), the combined organic phases are washed 3 times with 5% aqueous solution of ammonia (100 mL) and finally with water until a pH of 8 is reached. After drying with anhydrous sodium sulfate, the solvent of the organic phase is removed under vacuum and the residue is purified by column chromatography (AcOEt/*n*-Heptan 1/1) affording the target compound as a white solid (2.37g, 7.17mmol, 96.9% yield).

$C_{19}H_{23}FN_2O_2$ (330.40g.mol$^{-1}$), m.p. 118-121°C. $^1$H NMR ($CDCl_3$): 7.23-7.27 (m, 1$H_{arom}$), 6.88-7.01 (m, 2$H_{arom}$), 5.08-5.15 (m, 1H), 4.72-4.84 (m, 2H), 4.46 (d, *J*= 16Hz, 1H), 4.07 (d, *J*= 16Hz, 1H), 2.74-3.02 (m, 4H), 2.42-2.50 (m, 1H), 1.69-1.86 (m, 5H), 1.48-1.60 (m, 2H), 1.23-1.32 (m, 3H). $^{13}$C NMR ($CDCl_3$): 174.8 (s, 1C=O), 164.4 (s, 1C=O), 161.1 (d, *J*=247Hz, 1q$C_{arom}$), 137.2 (d, *J*=8Hz, 1q$C_{arom}$), 128.6 (s, 1q$C_{arom}$), 127.2 (d, *J*=8Hz, 1$C_{arom}$), 115.7 (d, *J*=21Hz, 1$C_{arom}$), 114.3 (d, *J*=21Hz, 1$C_{arom}$), 54.6 (s, 1CH), 49.0 (s, 1$CH_2$), 45.1 (s, 1$CH_2$), 38.9 (s, 1CH), 29.23 (s, 2$CH_2$), 29.0 (s, 1$CH_2$), 28.8 (s, 1$CH_2$), 25.7 (s, 3$CH_2$). MS (EI): m/z (%): 330 (59) [M]$^+$, 219 (98), 203 (33), 191 (20), 181 (12), 164 (39), 150 (100), 135 (9), 113 (24), 101 (4), 83 (41), 67 (4), 55 (51), 42 (28), 28 (6).

**(*R*)-2-(1-Chloro-cyclohexanecarbonyl)-1,2,3,6,7,11b-hexahydro-pyrazino[2,1-a]isoquinolin-4-one (15)**

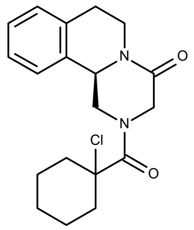

4g (*R*)-Praziquanamine#(19.78mmol) are dissolved in a mixture of DCM (100mL) and water (100mL) under nitrogen. Sodium hydroxide is added to reach a pH of 10. A solution of 3.96g 1-Chloro-cyclohexanecarbonyl chloride (20.77mmol) in DCM (50mL) and 2.6g of a 32% aqueous solution of sodium hydroxide (20.77mmol) are added dropwise in parallel to the biphasic mixture keeping the pH between 9 and 11. After addition the reaction mixture is stirred for 3h at RT and finally allowed to settle. The two phases are separated, the aqueous one is extracted 3 times with DCM (30mL), the combined organic phases are washed 3 times with a 5% aqueous solution of ammonia (50 mL) and then 3 times with water until a pH of 9 is reached. After drying with anhydrous sodium sulfate, the solvent is removed under vacuum. The crude residue is purified by column chromatography (AcOEt/*n*-Heptan 1/1) to afford the target compound as white solid (6.51g, 18.77mmol, 94.9% yield).##

$C_{19}H_{23}ClN_2O_2$ (346.85g.mol$^{-1}$), m.p. 135-137°C. $^1$H NMR ($CDCl_3$): 7.24-7.29 (m, 3$H_{arom}$), 7.18-7.21 (m, 1$H_{arom}$), 5.21 (d, *J*=15Hz, 1H), 4.94-5.10 (m, 2H), 4.86 (d, *J*= 15Hz, 1H), 2.76-3.05 (m, 4H), 2.02-2.21 (m, 4H), 1.76-1.87 (m, 2H), 1.63-1.71 (m, 4H), 1.24-1.36 (m, 1H). $^{13}$C NMR ($CDCl_3$): 169.4 (s, 1C=O), 164.5 (s, 1C=O), 135.1 (s, 1q$C_{arom}$), 132.5 (s, 1q$C_{arom}$), 129.5 (s, 1$C_{arom}$), 127.5 (s, 1$C_{arom}$), 127.0 (s, 1$C_{arom}$), 125.4 (s, 1$C_{arom}$), 70.7 (s, 1qC), 54.9 (s, 1CH), 38.8 (s, 1$CH_2$), 37.2 (s, 1$CH_2$), 28.8 (s, 2$CH_2$), 24.8 (s, 2$CH_2$), 22.0 (s, 2$CH_2$), 21.9 (s, 1$CH_2$). MS (EI): m/z (%): 346 (58) $[M]^+$, 201 (43), 185 (49), 180 (45), 146 (52), 132 (100), 81 (30), 42 (4).

**(*R*)-2-(4-Chloro-cyclohexanecarbonyl)-1,2,3,6,7,11b-hexahydro-pyrazino[2,1-a]isoquinolin-4-one (14)**

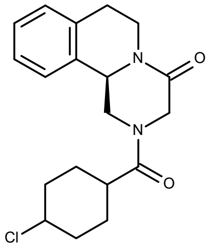


Same procedure starting from 4g (*R*)-Praziquanamine (19.78mmol) and 3.76g 4-Chloro-cyclohexanecarbonyl chloride (20.77mmol) affording after purification by column chromatography (EtOAc/*n*-heptane 1/1) the target compound as a white solid (6.64g, 19.14mmol, 96.8% yield).

$C_{19}H_{23}ClN_2O_2$ (346.85g.mol$^{-1}$). $^1$H NMR ($CDCl_3$): 7.23-7.30 (m, 3$H_{arom}$), 7.17-7.21 (m, 1$H_{arom}$), 5.13-5.18 (m, 1H), 4.79-4.86 (m, 2H), 4.40-4.52 (m, 1H), 3.82-3.94 (m, 1H), 2.78-3.02 (m, 4H), 2.22-2.35 (m, 1H), 2.02-2.17 (m, 2H), 1.78-1.93 (m, 2H) 1.58-1.72 (m, 5H). $^{13}$C NMR ($CDCl_3$): 165.3 (s, 1C=O), 153.8 (s, 1C=O), 135.0 (s, 1q$C_{arom}$), 132.8 (s, 1q$C_{arom}$), 129.3 (s, 1$C_{arom}$), 127.5 (s, 1$C_{arom}$), 127.0 (s, 1$C_{arom}$), 125.4 (s, 1$C_{arom}$), 58.1 (s, 1CH), 56.0 (s, 1qC), 54.9 (s, 1CH), 49.0 (s, 1$CH_2$), 45.3 (s, 1$CH_2$), 39.7 (s, 1CH), 39.0 (s, 1$CH_2$), 36.5 (s, 1$CH_2$), 36.1 (s, 1$CH_2$), 33.2 (s, 1$CH_2$), 28.7 (s, 1$CH_2$), 23.3 (s, 1$CH_2$). MS (EI): m/z (%): 346 (31) $[M]^+$, 310 (6), 215 (3), 201 (72), 185 (36), 173 (14), 146 (44), 132 (100), 117 (12), 103 (6), 81 (45), 55 (6), 42 (17).

**Synthesis of (*R*)-Praziquantel homologues via coupling reaction of (*R*)-Praziquanamine with carboxylic acids :**

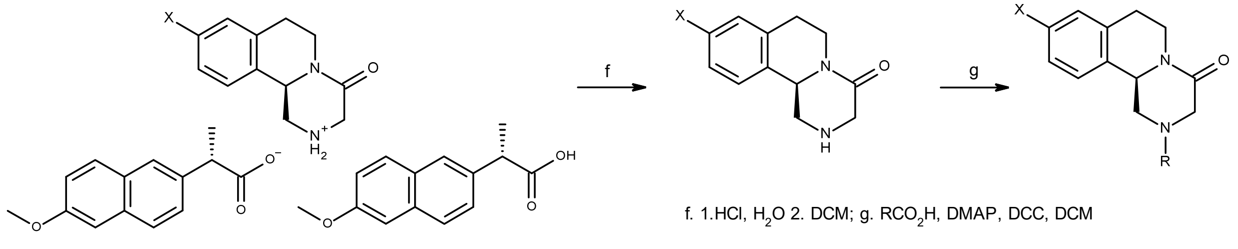


**(*R*)-9-Fluoro-2-(tetrahydro-pyran-4-carbonyl)-1,2,3,6,7,11b-hexahydro-pyrazino[2,1-a]isoquinolin-4-one (32)**

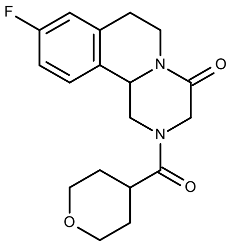


0.97g racemic fluorinated Praziquanamine (9-Fluoro-1,2,3,6,7,11b-hexahydro-pyrazino[2,1-a]isoquinolin-4-one) (4.40mmol), 0.16g DMAP (Dimethyl-pyridin-4-yl-amine) (1.32mmol) and 0.65g tetrahydro-pyran-4-carboxylic acid (4.84mmol) are dissolved in DCM (150mL) under nitrogen. A solution of 1.19g DCC (*N*,*N*'-Dicyclohexylcarbodiimide) (5.73mmol) in DCM (50mL) is added dropwise over 30min. The resulting mixture is stirred overnight at RT before addition of an aqueous solution of 0.33g oxalic acid (2.64 mmol) dissolved in water (40mL). After stirring for 30min, the white suspension was filtered and the filter cake washed 3 times with DCM (50mL). The phases are separated, the organic layer is washed twice with saturated solution of potassium carbonate (50mL) and finally water. After drying on anhydrous sodium sulfate, the solvent is removed under vacuum. The yellow residue is purified by column chromatography (AcOEt/*n*-Heptan 2/3) affording the target compound as a slightly yellow solid (0.88g, 2.65mmol, 60.2% yield).

$C_{18}H_{21}FN_2O_3$ (332.37g.mol$^{-1}$), m.p. 177-181°C. $^1$H NMR ($CDCl_3$): 7.21-7.28 (m, 1$H_{arom}$), 6.86-7.03 (m, 2$H_{arom}$), 5.05-5.17 (m, 1H), 4.72-4.84 (m, 2H), 4.45 (d, *J*= 18Hz, 1H), 4.09 (d, *J*= 18Hz, 1H), 3.99-4.04 (m, 2H), 3.40-3.52 (m, 2H), 2.68-3.02 (m, 4H), 1.82-2.02 (m, 2H), 1.58-1.71 (m, 3H). $^{13}$C NMR ($CDCl_3$): 173.0 (s, 1C=O), 164.0 (s, 1C=O), 161.7 (d, *J*=248Hz, 1q$C_{arom}$), 137.1 (d, *J*=7Hz, 1q$C_{arom}$), 128.4 (s, 1q$C_{arom}$), 127.1 (d, *J*=7Hz, 1$C_{arom}$), 115.7 (d, *J*=21Hz, 1$C_{arom}$), 114.3 (d, *J*=21Hz, 1$C_{arom}$), 67.1 (s, 2$CH_2$), 54.6 (s, 1CH), 49.0 (s, 1$CH_2$), 45.3 (s, 1$CH_2$), 38.9 (s, 1$CH_2$), 37.9 (s, 1CH), 28.8 (s, 1$CH_2$), 28.8 (s, 1$CH_2$), 28.7 (s, 1$CH_2$). MS (EI): m/z (%): 332 (68) $[M]^+$, 275 (4), 219 (62), 203 (26), 191 (15), 183 (5), 164 (39), 150 (100), 126 (11), 113 (18), 85 (9), 67 (8), 55 (47), 42 (28), 28 (18).

**(*R*)-2-(4-trans-methyl-cyclohexanecarbonyl)-1,2,3,6,7,11b-hexahydro-pyrazino[2,1-a]isoquinolin-4-one (7)**

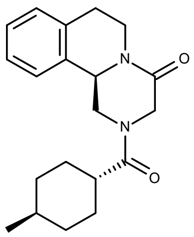

Same procedure starting from 10g (*R*)-Praziquanamine (49mmol) and 7.16g trans-4-methyl-cyclohexane carboxylic acid (49mmol) affording after purification by column chromatography (EtOAc/*n*-heptane 2/3 and 1/2) the target compound as a white solid (14.8g, 45.3mmol, 92.5% yield).

$C_{20}H_{26}N_2O_2$ (326.43 g.mol$^{-1}$), m.p. 90-93°C. $^1$H NMR ($CDCl_3$): 7.18-7.34 (m, 4$H_{arom}$), 5.19 (d, *J*=16Hz, 1H), 4.80-4.88 (m, 2H), 4.50 (d, *J*=17Hz, 1H), 4.11 (d, *J*=17Hz, 1H), 2.87-3.07 (m, 2H), 2.78-2.86 (m, 2H), 2.40-2.48 (m, 1H), 1.75-1.87 (m, 3H), 1.58-1.68 (m, 2H), 1.42-1.50 (m, 1H), 0.92-1.02 (m, 6H). $^{13}$C NMR ($CDCl_3$): 174.7 (s, 1C=O), 164.3 (s, 1C=O), 134.7 (s, 1q$C_{arom}$), 132.8 (s, 1q$C_{arom}$), 129.2 (s, 1$C_{arom}$), 127.4 (s, 1$C_{arom}$), 126.9 (s, 1$C_{arom}$), 125.4 (s, 1$C_{arom}$), 54.9 (s, 1CH), 49.0 (s, 1$CH_2$), 45.1 (s, 1$CH_2$), 40.5 (s, 1CH), 39.0 (s, 1$CH_2$), 34.3 (s, 2$CH_2$), 32.0 (s, 1CH), 29.2 (s, 1$CH_2$), 28.9 (s, 1$CH_2$), 28.7 (s, 1$CH_2$), 22.5 (s, 1$CH_3$). MS (EI) m/z (%): 326 (50) [M]$^+$, 201 (100), 185 (35), 173 (19), 146 (40), 132 (100), 97 (20), 55 (29), 42 (10).

**(*R*)-2-(4,4'-dimethyl-cyclohexanecarbonyl)-1,2,3,6,7,11b-hexahydro-pyrazino[2,1-a]isoquinolin-4-one (9)**

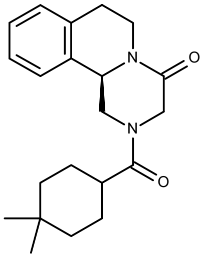

Same procedure starting from 5.75g (*R*)-Praziquanamine (28.43mmol) and 4.68g 4,4-dimethyl-cyclohexane carboxylic acid (28.46mmol) affording after purification by column chromatography (EtOAc/*n*-heptane 1/2, 1/1 and 4/3) the target compound as a white solid (6.89g, 20.24mmol, 71.2% yield).

$C_{21}H_{28}N_2O_2$ (340.46 g.mol$^{-1}$), m.p. 105-108°C. $^1$H NMR ($CDCl_3$): 7.20-7.33 (m, 4$H_{arom}$), 5.19 (d, *J*=16Hz, 1H), 4.81-4.89 (m, 2H), 4.48 (d, *J*=17Hz, 1H), 4.10 (d, *J*=17Hz, 1H), 2.88-3.06 (m, 2H), 2.78-2.86 (m, 2H), 2.36-2.43 (m, 1H), 1.72-1.85 (m, 3H), 1.57-1.65 (m, 2H), 1.48-1.56 (m, 2H), 1.22-1.31 (m, 1H), 0.98 (s, 3H), 0.96 (s, 6H). $^{13}$C NMR ($CDCl_3$): 174.8 (s, 1C=O), 164.4 (s, 1C=O), 134.8 (s, 1q$C_{arom}$), 132.8 (s, 1q$C_{arom}$), 129.3 (s, 1$C_{arom}$), 127.5 (s, 1$C_{arom}$), 127.0 (s, 1$C_{arom}$), 125.5 (s, 1$C_{arom}$), 55.0 (s, 1CH), 49.0 (s, 1$CH_2$), 45.2 (s, 1$CH_2$), 40.9 (s, 1CH), 39.1 (s, 1$CH_2$), 38.5 (s, 2$CH_2$), 32.8 (s, 1$CH_3$), 29.9 (s, 1qC), 28.7 (s, 1$CH_2$), 25.2 (s, 1$CH_2$), 25.0 (s, 1$CH_2$), 24.2 (s, 1$CH_3$). MS (EI) m/z (%): 340 (45) [M]$^+$, 201 (100), 185 (34), 173 (20), 146 (40), 132 (100), 113 (18), 69 (32), 55 (19), 42 (13).

**(*R*)-2-(Bicyclo[3.1.0]hexane-3-carbonyl)-1,2,3,6,7,11b-hexahydro-pyrazino[2,1-a]isoquinolin-4-one (6)**

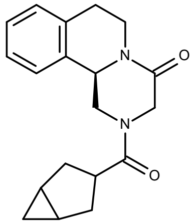

Same procedure starting from 8.27g (*R*)-Praziquanamine (40.90mmol) and 5.16g bicyclo[3.1.0]hexane-3-carboxylic acid (40.90mmol) affording after crystallization from *n*-heptane/ethanol the target compound as a white solid (9.84g, 31.70mmol, 77.5% yield).

$C_{19}H_{22}N_2O_2$ (310.40g.mol$^{-1}$), m.p. 147-150°C. $^1$H NMR ($CDCl_3$): 7.16-7.26 (m, 4$H_{arom}$), 5.11 (dt, *J*=15, 3Hz, 1H), 4.76-4.88 (m, 2H), 4.36 (d, *J*=18Hz, 1H), 4.00 (d, *J*=18Hz, 1H), 3.12-3.27 (m, 2H), 2.75-3.03 (m, 4H), 2.17-2.33 (m, 2H), 1.95-1.99 (m, 1H), 1.29-1.39 (m, 2H), 0.45-0.52 (m, 1H), 0.30-0.35 (m, 1H). $^{13}$C NMR ($CDCl_3$): 175.3 (s, 1C=O), 164.2 (s, 1C=O), 134.7 (s, 1q$C_{arom}$), 132.8 (s, 1q$C_{arom}$), 129.3 (s, 1$C_{arom}$), 127.4 (s, 1$C_{arom}$), 126.9 (s, 1$C_{arom}$), 125.4 (s, 1$C_{arom}$), 54.9 (s, 1CH), 49.3 (s, 1$CH_2$), 45.6 (s, 1$CH_2$), 41.3 (s, 1CH), 39.1 (s, 1$CH_2$), 31.9 (s, 1$CH_2$), 28.7 (s, 1$CH_2$), 18.6 (s, 1CH), 18.5 (s, 1CH), 10.9 (s, 2$CH_2$). MS (EI) m/z (%): 310 (29) [M]$^+$, 201 (48), 185 (14), 173 (14), 145 (33), 132 (100), 117 (10), 81 (23), 42 (18).

**(*R*)-2-(Bicyclo[2.2.2]octane-1-carbonyl)-1,2,3,6,7,11b-hexyhadro-pyrazino[2,1-a]isoquinolin-4-one (12)**

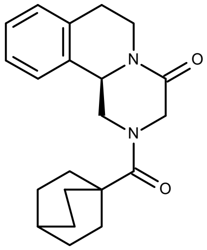

Same procedure starting from 6.95g (*R*)-Praziquanamine (34.37mmol) and 7.88g bicyclo[2.2.2]octane-1-carboxylic acid (34.37mmol) affording after crystallization from *n*-heptane/ethanol the target compound as a white solid (8.41g, 24.85mmol, 72.3% yield).

$C_{21}H_{26}N_2O_2$ (338.45g.mol$^{-1}$), m.p. 167-170°C. $^1$H NMR ($CDCl_3$): 7.16-7.31 (m, 4$H_{arom}$), 5.12 (dt, *J*=15, 3Hz, 1H), 4.78-4.89 (m, 3H), 3.96 (d, *J*=18Hz, 1H), 2.73-3.03 (m, 4H), 1.86-1.91 (m, 6H), 1.60-1.71 (m, 7H). $^{13}$C NMR ($CDCl_3$): 176.1 (s, 1C=O), 164.7 (s, 1C=O), 134.9 (s, 1q$C_{arom}$), 132.9 (s, 1q$C_{arom}$), 129.4 (s, 1$C_{arom}$), 127.4 (s, 1$C_{arom}$), 126.9 (s, 1$C_{arom}$), 125.2 (s, 1$C_{arom}$), 54.9 (s, 1CH), 50.5 (s, 1$CH_2$), 48.3 (s, 1$CH_2$), 39.5 (s, 1qC), 39.1 (s, 1$CH_2$), 28.8 (s, 1$CH_2$), 28.1 (s, 3$CH_2$), 25.4 (s, 3$CH_2$), 23.7 (s, 1CH). MS (EI) m/z (%): 338 (27) [M]$^+$, 201 (100), 185 (26), 146 (43), 132 (85), 109 (66), 67 (55), 42 (13).

**(*R*)-2-(Spiro[2.5]octane-6-carbonyl)-1,2,3,6,7,11b-hexahydro-pyrazino[2,1-a]isoquinolin-4-one (10)**

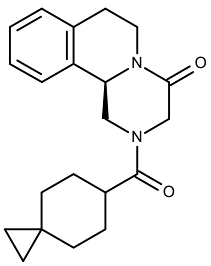


Same procedure starting from 2.39g (*R*)-Praziquanamine (11.79mmol) and 1.82g spiro[2.5]octane-6-carboxylic acid (11.79mmol) affording after column chromatography (EtOAc/*n*-heptane 3/7 and 3/2) the target compound as a white solid (3.65g, 10.78mmol, 91.4% yield).

$C_{21}H_{26}N_2O_2$ (338.45g.mol$^{-1}$), m.p. 155-158°C. $^1$H NMR ($CDCl_3$): 7.16-7.27 (m, 4$H_{arom}$), 5.16 (dd, *J*=12, 4Hz, 1H), 4.77-4.86 (m, 3H), 4.47 (d, *J*=16Hz, 1H), 4.08 (d, *J*=16Hz, 1H), 2.75-3.01 (m, 3H), 2.46-2.50 (m, 1H), 1.73-1.81 (m, 5H), 0.93-0.99 (m, 2H), 0.21-0.32 (m, 5H). $^{13}$C NMR ($CDCl_3$): 174.7 (s, 1C=O), 164.4 (s, 1C=O), 134.8 (s, 1q$C_{arom}$), 132.8 (s, 1q$C_{arom}$), 129.3 (s, 1$C_{arom}$), 127.5 (s, 1$C_{arom}$), 127.0 (s, 1$C_{arom}$), 125.5 (s, 1$C_{arom}$), 54.9 (s, 1CH), 49.1 (s, 1$CH_2$), 45.3 (s, 1$CH_2$), 40.4 (s, 1CH), 39.1 (s, 1$CH_2$), 34.7 (s, 1$CH_2$), 28.8 (s, 1$CH_2$), 28.3 (s, 1$CH_2$), 28.1 (s, 1$CH_2$), 18.3 (s, 1qC), 18.0 (s, 1$CH_2$), 12.4 (s, 1$CH_2$), 11.9 (s, 1$CH_2$). MS (EI) m/z (%): 338 (22) [M]$^+$, 201 (70), 146 (34), 132 (100), 67 (39), 55 (21), 42 (23).

**(*R*)-2-Cyclopentanecarbonyl-1,2,3,6,7,11b-hexahydro-pyrazino[2,1-a]isoquinolin-4-one (5)**

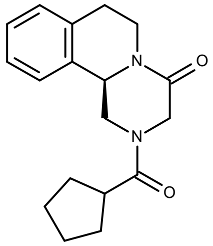


Same procedure starting from 17.70g (*R*)-Praziquanamine (87.61mmol) and 10g cyclopentane carboxylic acid (87.61mmol) affording after crystallization from n-heptane/ethanol the target compound as a white solid (16.04g, 53.76mmol, 61.4% yield).

$C_{18}H_{22}N_2O_2$ (298.39g.mol$^{-1}$), m.p. 137-139°C. $^1$H NMR (DMSO-$d_6$): 7.47-7.49 (m, 1$H_{arom}$), 7.20-7.28 (m, 3$H_{arom}$), 4.95 (dd, *J*=10, 5Hz, 1H), 4.79-4.85 (m, 1H), 4.51-4.59 (m, 2H), 4.41 (d, *J*=15Hz, 1H), 3.71 (d, *J*=15Hz, 1H), 3.23-3.28 (m, 1H), 2.99-3.05 (m, 1H), 2.74-2.95 (m, 3H), 1.74-1.82 (m, 2H), 1.52-1.70 (m, 5H). $^{13}$C NMR (DMSO-$d_6$): 173.8 (s, 1C=O), 164.6 (s, 1C=O), 135.0 (s, 1q$C_{arom}$), 133.2 (s, 1q$C_{arom}$), 129.0 (s, 1$C_{arom}$), 127.0 (s, 1$C_{arom}$), 126.6 (s, 1$C_{arom}$), 125.8 (s, 1$C_{arom}$), 54.8 (s, 1CH), 54.0 (s, 1CH), 48.5 (s, 1$CH_2$), 48.1 (s, 1$CH_2$), 45.8 (s, 1$CH_2$), 44.6 (s, 1$CH_2$), 29.6 (s, 1$CH_2$), 29.4 (s, 1$CH_2$), 28.2 (s, 1$CH_2$), 25.7 (s, 1$CH_2$). MS (EI) m/z (%): 298 (27) [M]$^+$, 201 (48), 185 (21), 146 (31), 132 (100), 126 (19), 69 (44), 49 (18), 41 (31).

**(*R*)-2-(4-Methylene-cyclohexanecarbonyl)-1,2,3,6,7,11b-hexahydro-pyrazino[2,1-a]isoquinolin-4-one (8)**

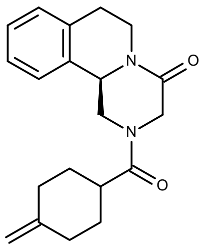

Same procedure starting from 2.78g (*R*)-Praziquanamine (13.77mmol) and 1.93g 4-methylene-cyclohexane carboxylic acid (13.77mmol) affording after column chromatography (EtOAc/*n*-heptane 2/3 and 3/2) the target compound as a white solid (3.78g, 11.65mmol, 85% yield).

$C_{20}H_{24}N_2O_2$ (324.42g.mol$^{-1}$), decomposition at 105-108°C. $^1$H NMR (DMSO-d$_6$): 7.17-7.28 (m, 4H$_{arom}$), 5.16 (dd, *J*=15, 5Hz, 1H), 4.76-4.91 (m, 3H), 4.68 (s, 2H), 4.49 (d, *J*=20Hz, 1H), 4.07-4.14 (m, 1H), 2.77-3.02 (m, 4H), 2.59-2.64 (m, 1H), 2.38-2.43 (m, 2H), 2.05-2.11 (m, 2H), 1.59-1.71 (m, 2H), 1.24-1.27 (m, 1H). $^{13}$C NMR (DMSO-d$_6$): 173.3 (s, 1C=O), 164.6 (s, 1C=O), 134.9 (s, 1qC$_{arom}$), 133.2 (s, 1qC$_{arom}$), 128.9 (s, 1C$_{arom}$), 127.5 (s, 1C$_{arom}$), 126.9 (s, 1C$_{arom}$), 125.2 (s, 1C$_{arom}$), 125.1 (s, 1qC), 108.3 (s, 1CH$_2$), 54.8 (s, 1CH), 53.9 (s, 1CH$_2$), 48.4 (s, 1CH$_2$), 45.7 (s, 1CH$_2$), 38.5 (s, 1CH$_2$), 38.0 (s, 1CH$_2$), 33.1 (s, 1CH$_2$), 30.3 (s, 1CH$_2$), 30.0 (s, 1CH$_2$), 28.1 (s, 1CH). MS (EI) m/z (%): 324 (26) [M]$^+$, 201 (57), 146 (27), 132 (100), 95 (28), 67 (20), 55 (21), 42 (21).

**3-((*R*)-4-Oxo-1,3,4,6,7,11b-hexahydro-pyrazino[2,1-a]isoquinoline-2-carbonyl)-piperidine-1-carboxylic acid *tert*-butyl ester**

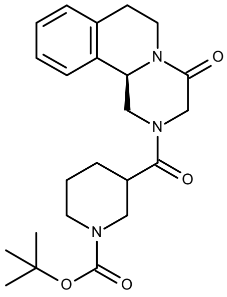

Same procedure starting from 8.30g (*R*)-Praziquanamine (41mmol) and 9.58g piperidine-1,3-dicarboxylic acid 1-tert-butyl ester (41mmol) affording after column chromatography (EtOAc/*n*-heptane 2/3 and 1/2) the target compound as a white solid (14.57g, 35mmol, 86% yield).

$C_{23}H_{31}N_3O_4$ (413.51 g.mol$^{-1}$), m.p. 162-165°C. $^1$H NMR (CDCl$_3$): 7.19-7.33 (m, 4H$_{arom}$), 5.08-5.22 (m, 1H), 4.77-4.93 (m, 3H), 4.44-4.54 (m, 1H), 4.14 (t, *J*=16Hz, 3H), 2.62-3.06 (m, 8H), 1.88-2.02 (m, 2H), 1.51 (s, 9H). $^{13}$C NMR (CDCl$_3$): 172.0 (s, 1C=O), 165.1 (s, 1C=O), 164.0 (s, 1C=O), 134.7 (s, 1qC$_{arom}$), 132.6 (s, 1qC$_{arom}$), 129.4 (s, 1C$_{arom}$), 127.5 (s, 1C$_{arom}$), 127.0 (s, 1C$_{arom}$), 125.4 (s, 1C$_{arom}$), 79.9 (s, 1qC), 54.9 (s, 1CH), 49.1 (s, 1CH), 49.0 (s, 1CH$_2$), 46.3 (s, 1CH$_2$), 45.3 (s, 1CH$_2$), 39.1 (s, 2CH$_2$), 28.7 (s, 2CH$_2$), 28.5 (s, 3CH$_3$), 27.6 (s, 1CH$_2$). MS (EI) m/z (%): 413 (4) [M]$^+$, 357 (60), 340 (11), 312 (47), 257 (3), 229 (6), 201 (64), 173 (24), 146 (100), 132 (76), 110 (58), 84 (82), 57 (92), 49 (28), 41 (25).

**(*R*)-2-(piperidine-3-carbonyl)-1,2,3,6,7,11b-hexahydro-pyrazino[2,1-a]isoquinolin-4-one (29)**

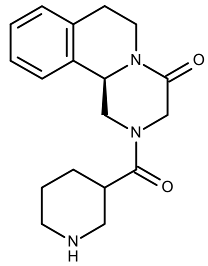

4g *N*-Boc-protected compound (3-((R)-4-Oxo-1,3,4,6,7,11b-hexahydro-pyrazino[2,1-a]isoquinoline-2-carbonyl)-piperidine-1-carboxylic acid *tert*-butyl ester) (9.67mmol) are suspended in toluene (80mL) and stirred at RT under nitrogen. 11.7mL trifluoroacetic acid (152mmol) are added dropwise, the suspension dissolving slowly. After stirring for 6h, 14mL of a 32% aqueous solution of sodium hydroxide (152mmol) are added, the phases are subsequently separated, the aqueous layer is extracted 5 times with toluene (60mL). The combined organic phases are dried over sodium sulfate. Isopropanol (60mL) is added and the solvent is removed under vacuum. The residue is finally purified by column chromatography (DCM/MeOH 4/1, 7/3 and 7/4) affording the target compound as slightly yellow solid (2.33g, 7.43mmol, 76.9% yield).

$C_{18}H_{23}N_3O_2$ (313.40 g.mol$^{-1}$), m.p. 155-162°C. $^1$H NMR ($CDCl_3$): 7.15-7.26 (m, 4H$_{arom}$), 5.07-5.13 (m, 1H), 4.73-4.80 (m, 2H), 4.43 (d, *J*=16Hz, 1H), 4.06 (d, *J*=16Hz, 1H), 2.62-3.15 (m, 8H), 2.61-2.68 (m, 2H), 2.59-2.77 (m, 1H), 1.85 (s, 2H), 1.47-1.52 (m, 1H). $^{13}$C NMR ($CDCl_3$): 173.1 (s, 1C=O), 164.2 (s, 1C=O), 134.7 (s, 1qC$_{arom}$), 132.7 (s, 1qC$_{arom}$), 129.3 (s, 1C$_{arom}$), 127.5 (s, 1C$_{arom}$), 127.0 (s, 1C$_{arom}$), 125.5 (s, 1C$_{arom}$), 55.8 (s, 1CH), 54.9 (s, 1CH), 48.9 (s, 1CH$_2$), 46.4 (s, 1CH$_2$), 45.1 (s, 1CH$_2$), 39.1 (s, 1CH$_2$), 28.7 (s, 1CH$_2$), 28.0 (s, 1CH$_2$), 27.6 (s, 1CH$_2$), 25.6 (s, 1CH$_2$). MS (EI) m/z (%): 313 (46) [M]$^+$, 257 (4), 201 (18), 173 (15), 145 (25), 132 (39), 84 (100), 68 (8), 56 (15), 42 (12), 28 (4).

**2-((*R*)-4-Oxo-1,3,4,6,7,11b-hexahydro-pyrazino[2,1-a]isoquinoline-2-carbonyl)-piperidine-1-carboxylic acid *tert*-butyl ester**

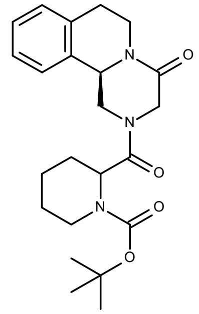

Same procedure starting from 10g (*R*)-Praziquanamine (49mmol) and 11.54g piperidine-1,2-dicarboxylic acid 1-tert-butyl ester (49mmol) affording after column chromatography (EtOAc/*n*-heptane 3/2 and 2/1) the target compound as a white solid (17.67g, 43mmol, 87% yield).

$C_{23}H_{31}N_3O_4$ (413.51 g.mol$^{-1}$), m.p. 104-111°C. $^1$H NMR ($CDCl_3$): 7.19-7.29 (m, 4H$_{arom}$), 5.05-5.22 (m, 1H), 4.78-4.96 (m, 3H), 4.41-4.52 (m, 1H), 4.06-4.15 (m, 1H), 3.89-3.98 (m, 1H), 3.73-3.82 (m, 1H), 3.08-3.19 (m, 1H), 2.74-3.03 (m, 4H), 1.90-2.04 (m, 1H), 1.61-1.77 (m, 4H), 1.47 (s, 9H). $^{13}$C NMR ($CDCl_3$): 171.1 (s, 1C=O), 165.2 (s, 1C=O), 164.0 (s, 1C=O), 135.0 (s, 1qC$_{arom}$), 132.7 (s, 1qC$_{arom}$), 129.3 (s, 1C$_{arom}$), 127.5 (s, 1C$_{arom}$), 126.9 (s, 1C$_{arom}$), 125.5 (s, 1C$_{arom}$), 80.6 (s, 1qC), 55.7 (s, 1CH), 55.0 (s, 1CH), 49.1 (s, 1CH$_2$), 46.7 (s, 1CH$_2$), 42.7 (s, 1CH$_2$), 39.1 (s, 1CH$_2$), 38.8 (s, 1CH$_2$), 34.0 (s, 1CH$_2$), 28.8 (s, 1CH$_2$), 28.4 (s, 3CH$_3$), 26.5 (s, 1CH$_2$). MS (EI) m/z (%): 413 (1) [M]$^+$, 357 (2), 340 (3), 312 (3), 257 (2), 229 (2), 201 (5), 184 (20), 173 (5), 145 (6), 128 (100), 117 (5), 84 (86), 57 (25), 49 (5), 41 (6).

**(*R*)-2-(Piperidine-2-carbonyl)-1,2,3,6,7,11b-hexahydro-pyrazino[2,1-α]isoquinolin-4-one (28)**

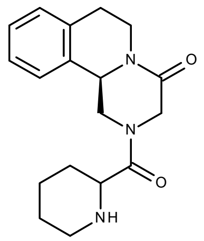


$C_{18}H_{23}N_3O_2$ (313.40 g.mol$^{-1}$), m.p. 115-121°C. $^1$H NMR ($CDCl_3$): 7.16-7.29 (m, 4$H_{arom}$), 5.12-5.17 (m, 1H), 4.76-4.86 (m, 2H), 4.67 (d, *J*=16Hz, 1H), 4.04 (d, *J*=16Hz, 1H), 3.52-3.62 (m, 1H), 3.12-3.21 (m, 1H), 2.83-3.02 (m, 3H), 2.62-2.78 (m, 2H), 1.91-1.97 (m, 1H), 1.71-1.79 (m, 1H), 1.45-1.62 (m, 3H), 1.32-1.43 (m, 2H). $^{13}$C NMR ($CDCl_3$): 172.1 (s, 1C=O), 164.2 (s, 1C=O), 134.8 (s, 1q$C_{arom}$), 132.6 (s, 1q$C_{arom}$), 129.3 (s, 1$C_{arom}$), 127.5 (s, 1$C_{arom}$), 127.0 (s, 1$C_{arom}$), 125.5 (s, 1$C_{arom}$), 57.0 (s, 1CH), 54.8 (s, 1CH), 48.8 (s, 1$CH_2$), 46.0 (s, 1$CH_2$), 45.4 (s, 1$CH_2$), 39.1 (s, 1$CH_2$), 29.7 (s, 1$CH_2$), 28.7 (s, 1$CH_2$), 26.8 (s, 1$CH_2$), 24.3 (s, 1$CH_2$). MS (EI) m/z (%): 313 (1) $[M]^+$, 285 (2), 257 (1), 202 (2), 173 (3), 145 (6), 132 (10), 84 (100), 67 (3), 56 (10), 42 (5), 28 (3).

**(*R*)-2-(Tetrahydro-furan-2-carbonyl)-1,2,3,6,7,11b-hexahydro-pyrazino[2,1-a]isoquinolin-4-one (24)**

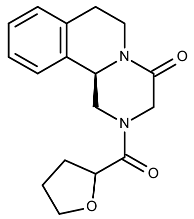


Same procedure starting from 10g (*R*)-Praziquanamine (49mmol) and 6.51g tetrahydrofuran-2-carboxylic acid (54.39mmol) affording after column chromatography (EtOAc/*n*-heptane 3/1) the target compound as a white solid (7.45g, 24.8mmol, 50% yield).

$C_{17}H_{20}N_2O_3$ (300.35 g.mol$^{-1}$), m.p. 120-124°C. $^1$H NMR ($CDCl_3$): 7.25-7.32 (m, 3$H_{arom}$), 7.19-7.23 (m, 1$H_{arom}$), 4.98-5.17 (m, 1H), 4.80-4.95 (m, 2H), 4.62-4.73 (m, 2H), 3.79-4.00 (m, 3H), 2.95-3.20 (m, 1H), 2.77-2.94 (m, 3H), 2.12-2.35 (m, 1H), 1.92-2.09 (m, 3H). $^{13}$C NMR ($CDCl_3$): 170.3 (s, 1C=O), 165.0 (s, 1C=O), 135.2 (s, 1q$C_{arom}$), 132.7 (s, 1q$C_{arom}$), 129.4 (s, 1$C_{arom}$), 127.4 (s, 1$C_{arom}$), 126.9 (s, 1$C_{arom}$), 125.5 (s, 1$C_{arom}$), 69.3 (s, 1$CH_2$), 56.1 (s, 1CH), 54.8 (s, 1CH), 49.5 (s, 1$CH_2$), 46.0 (s, 1$CH_2$), 38.8 (s, 1$CH_2$), 28.7 (s, 1$CH_2$), 27.4 (s, 1$CH_2$), 25.9 (s, 1$CH_2$). MS (EI) m/z (%): 300 (56) $[M]^+$, 272 (3), 257 (1), 229 (3), 201 (67), 185 (38), 173 (33), 145 (71), 132 (100), 117 (14), 103 (7), 91 (4), 77 (4), 71 (79), 56 (3), 43 (25), 27 (2).

**(*R*)-2-(Tetrahydro-thiopyran-4-carbonyl)-1,2,3,6,7,11b-hexahydro-pyrazino[2,1-a]isoquinolin-4-one (16)**

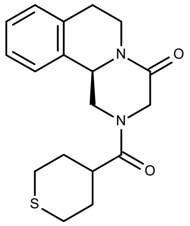


Same procedure starting from 6.82g (*R*)-Praziquanamine (33.72mmol) and 4.93g tetrahydro-thiopyran-4-carboxylic acid (33.72mmol) affording after column chromatography (EtOAc/*n*-heptane 3/2 and 4/1) the target compound as a white solid (9.39g, 28.42mmol, 84.3% yield).

$C_{18}H_{22}N_2O_2S$ (330.45 g.mol$^{-1}$), m.p. 146-149°C. $^{1}$H NMR ($CDCl_3$): 7.14-7.29 (m, 4$H_{arom}$), 5.06-5.10 (dd, *J*=12, 4Hz, 1H), 4.73-4.87 (m, 2H), 4.39 (d, *J*=16Hz, 1H), 4.05 (d, *J*=16Hz, 1H), 2.68-3.05 (m, 8H), 2.50-2.59 (m, 1H), 1.88-2.01 (m, 4H). $^{13}$C NMR ($CDCl_3$): 173.3 (s, 1C=O), 164.0 (s, 1C=O), 134.7 (s, 1q$C_{arom}$), 132.6 (s, 1q$C_{arom}$), 129.7 (s, 1$C_{arom}$), 127.8 (s, 1$C_{arom}$), 127.0 (s, 1$C_{arom}$), 125.4 (s, 1$C_{arom}$), 55.6 (s, 1CH), 49.6 (s, 1$CH_2$), 45.2 (s, 1$CH_2$), 40.2 (s, 1CH), 39.1 (s, 1$CH_2$), 30.1 (s, 1$CH_2$), 30.0 (s, 1$CH_2$), 28.7 (s, 1$CH_2$), 27.8 (s, 2$CH_2$). MS (EI) m/z (%): 330 (52) $[M]^+$, 201 (57), 185 (13), 173 (11), 146 (89), 132 (100), 117 (11), 101 (42), 67 (21), 55 (23).

**(*R*)-2-(Tetrahydro-thiopyran-3-carbonyl)-1,2,3,6,7,11b-hexahydro-pyrazino[2,1-a]isoquinolin-4-one (17)**

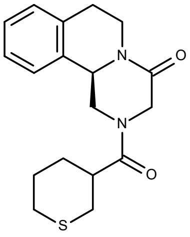


Same procedure starting from 6.07g (*R*)-Praziquanamine (30.02mmol) and 4.84g tetrahydro-thiopyran-3-carboxylic acid (33.02mmol) affording after column chromatography (EtOAc/*n*-heptane 3/2) the target compound as a white solid (8.61g, 26.06mmol, 86.8% yield).

$C_{18}H_{22}N_2O_2S$ (330.45 g.mol$^{-1}$), m.p. 107-110°C.$^{1}$H NMR ($CDCl_3$): 7.15-7.29 (m, 4$H_{arom}$), 5.06-5.15 (m, 1H), 4.73-4.89 (m, 3H), 4.43-4.52 (m, 1H), 4.10 (d, *J*=18Hz, 1H), 2.62-3.05 (m, 8H), 2.50-2.59 (m, 2H), 2.10-2.19 (m, 1H), 1.82-1.94 (m, 1H). $^{13}$C NMR ($CDCl_3$): 173.3 (s, 1C=O), 164.0 (s, 1C=O), 134.8 (s, 1q$C_{arom}$), 132.6 (s, 1q$C_{arom}$), 129.3 (s, 1$C_{arom}$), 127.5 (s, 1$C_{arom}$), 127.0 (s, 1$C_{arom}$), 125.4 (s, 1$C_{arom}$), 55.0 (s, 1CH), 49.0 (s, 1$CH_2$), 45.3 (s, 1$CH_2$), 41.7 (s, 1CH), 39.1 (s, 1$CH_2$), 30.4 (s, 1$CH_2$), 29.4 (s, 1$CH_2$), 28.7 (s, 1$CH_2$), 28.3 (s, 1$CH_2$), 27.3 (s, 2$CH_2$). MS (EI) m/z (%): 330 (40) $[M]^+$, 201 (64), 185 (11), 173 (15), 146 (79), 132 (100), 115 (14), 101 (37), 67 (29), 55 (24), 42 (34), 28 (51).

**(*R*)-2-(Tetrahydro-thiopyran-2-carbonyl)-1,2,3,6,7,11b-hexahydro-pyrazino[2,1-a]isoquinolin-4-one (18)**

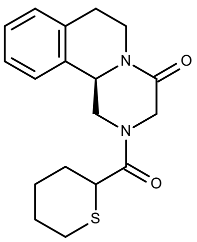


Same procedure starting from 6.07g (*R*)-Praziquanamine (30.02mmol) and 4.63g tetrahydro-thiopyran-2-carboxylic acid (31.50mmol) affording after column chromatography (EtOAc/*n*-heptane 3/2) the target compound as a white solid (7.21g, 21.82mmol, 72.7% yield).

$C_{18}H_{22}N_2O_2S$ (330.45 g.mol$^{-1}$), m.p. 107-110°C. $^1$H NMR ($CDCl_3$): 7.14-7.32 (m, 4$H_{arom}$), 5.02-5.18 (m, 1H), 4.75-4.93 (m, 2H), 4.45-4.68 (m, 1H), 4.05-4.28 (m, 1H), 3.52-3.68 (m, 1H), 2.71-3.02 (m, 6H), 1.90-2.15 (m, 5H), 1.67-1.85 (m, 1H). $^{13}$C NMR ($CDCl_3$): 170.0 (s, 1C=O), 164.2 (s, 1C=O), 135.3 (s, 1q$C_{arom}$), 132.1 (s, 1q$C_{arom}$), 129.3 (s, 1$C_{arom}$), 127.5 (s, 1$C_{arom}$), 127.0 (s, 1$C_{arom}$), 125.6 (s, 1$C_{arom}$), 54.8 (s, 1CH), 49.1 (s, 1$CH_2$), 45.6 (s, 1$CH_2$), 42.0 (s, 1CH), 39.0 (s, 1$CH_2$), 29.3 (s, 1$CH_2$), 28.8 (s, 1$CH_2$), 26.6 (s, 2$CH_2$), 24.7 (s, 1$CH_2$). MS (EI) m/z (%): 330 (27) [M]$^+$, 201 (29), 185 (16), 173 (18), 146 (100), 132 (86), 115 (16), 101 (73), 84 (14), 67 (27), 55 (18), 42 (38), 28 (69).

**(*R*)-2-(5-Oxo-tetrahydro-furan-2-carbonyl)-1,2,3,6,7,11b-hexahydro-pyrazino[2,1-a]isoquinolin-4-one (25)**

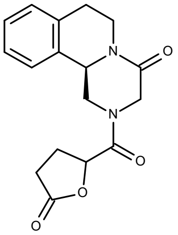


Same procedure starting from 6.88g (*R*)-Praziquanamine (34mmol) and 4.47g 5-oxo-tetrahydro-furan-2-carboxylic acid (34.34mmol) affording after column chromatography (EtOAc/*n*-heptane 3/1) the target compound as a white solid (9.15g, 29.11mmol, 85.6% yield).

$C_{17}H_{18}N_2O_4$ (314.34 g.mol$^{-1}$), m.p. 101-105°C. $^1$H NMR (DMSO-d6): 7.45-7.52 (m, 1$H_{arom}$), 7.21-7.32 (m, 3$H_{arom}$), 5.72-5.79 (m, 1H), 5.00-5.05 (m, 1H), 4.71-4-90 (m, 1H), 4.42-4.61 (m, 3H), 4.00-4.20 (m, 1H), 3.79-3.84 (m, 1H), 3.33-3.42 (m, 1H), 2.78-3.05 (m, 3H), 2.35-2.48 (m, 1H), 2.12-2.28 (m, 1H). $^{13}$C NMR (DMSO-d6): 177.4 (s, 1C=O), 168.0 (s, 1C=O), 164.4 (s, 1C=O), 135.6 (s, 1q$C_{arom}$), 133.3 (s, 1q$C_{arom}$), 129.5 (s, 1$C_{arom}$), 127.6 (s, 1$C_{arom}$), 127.0 (s, 1$C_{arom}$), 126.5 (s, 1$C_{arom}$), 74.5 (s, 1CH), 54.9 (s, 1CH), 48.2 (s, 1$CH_2$), 46.2 (s, 1$CH_2$), 38.8 (s, 1$CH_2$), 28.6 (s, 1$CH_2$), 27.2 (s, 1$CH_2$), 25.2 (s, 1$CH_2$). MS (EI) m/z (%): 314 (51) [M]$^+$, 200 (32), 185 (9), 173 (12), 155 (6), 145 (91), 132 (100), 117 (19), 103 (10), 85 (30), 77 (7), 57 (6), 42 (22), 29 (16).

**Synthesis of Praziquantel homologue :**

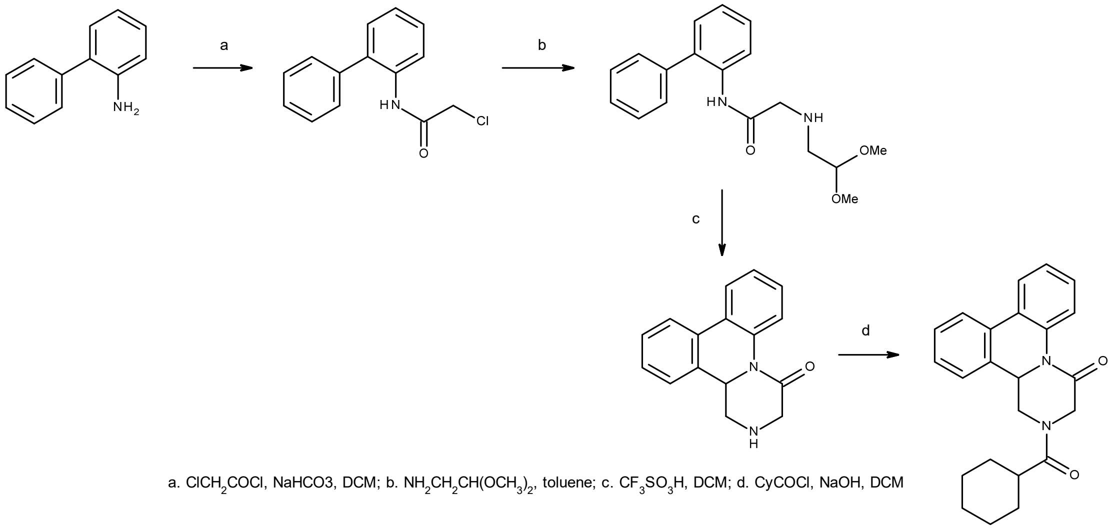


a. ClCH$_2$COCl, NaHCO3, DCM; b. NH$_2$CH$_2$CH(OCH$_3$)$_2$, toluene; c. CF$_3$SO$_3$H, DCM; d. CyCOCl, NaOH, DCM

**2-chloro-*N*-(2-phenylphenyl)acetamide**

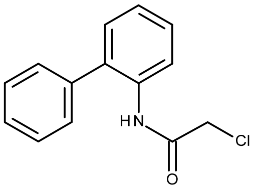


43.5g biphenyl-2ylamine (257.06mmol) and 32.39g sodium bicarbonate (385.59mmol) are dissolved in DCM (250mL). After cooling down the stirred solution at a temperature between -5°C and 0°C, 24.55mL chloro-acetyl chloride (308.47mmol) are slowly added dropwise under nitrogen. The reaction mixture is then stirred for 4h within a temperature range from -5°C to 5°C before addition of water (200 mL). After settling for 10 min, the phases are separated, the organic phase is washed with a 10% aqueous solution of sodium bicarbonate (150mL) and finally with water until the pH is in a range of 7. After drying over sodium sulfate, the solvent is removed under vacuum affording the target compound as a white solid (62.33g, 253.68mmol, 99% yield).

$C_{14}H_{12}ClNO$ (245.70 g.mol$^{-1}$), m.p. 96-98°C. $^{1}$H NMR (CDCl$_3$): 8.45 (bs, 1NH), 7.47-7.51 (m, 2H$_{arom}$), 7.37-7.45 (m, 5H$_{arom}$), 7.28-7.31 (m, 1H$_{arom}$), 7.21-7.25 (m, 1H$_{arom}$), 4.07 (s, 2H). $^{13}$C NMR (CDCl$_3$): 163.7 (s, 1C=O), 137.4 (s, 1qC$_{arom}$), 133.9 (s, 1qC$_{arom}$), 132.7 (s, 1qC$_{arom}$), 130.1 (s, 1C$_{arom}$), 129.3 (s, 2C$_{arom}$), 129.1 (s, 2C$_{arom}$), 128.5 (s, 1C$_{arom}$), 128.2 (s, 1C$_{arom}$), 125.0 (s, 1C$_{arom}$), 120.7 (s, 1C$_{arom}$), 43.0 (s, 1CH$_2$). MS (EI): m/z (%): 245 (96) [M]$^+$, 196 (59), 178 (46), 169 (100), 152 (16), 139 (18), 115 (12), 89 (6), 77 (11), 63 (6), 49 (8).

**2-(2,2-dimethoxyethylamino)-*N*-(2-phenylphenyl)acetamide**

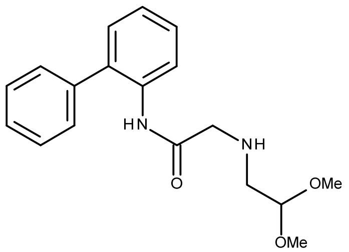


30g 2-chloro-N-(2-phenylphenyl)acetamide (122.10mmol) and 32.73g 2,2-Dimethoxy-ethylamine (311.35mmol) are dissolved in toluene (150mL) under nitrogen and the resulting solution is refluxed for 6h. The mixture is then cooled down, water (100mL) and toluene (100mL) are added and the phases are separated. The aqueous layer is extracted 3 times with toluene (100mL) and the combined organic layers are washed twice with water (80mL). The solvent is finally removed under vacuum affording the target compound as a yellow oil (35.71g, 113.59mmol, 93% yield).

$C_{18}H_{22}N_2O_3$ (314.38g.mol$^{-1}$). $^1$H NMR ($CDCl_3$): 9.45 (bs, 1NH), 8.47-8.50 (m, 1NH), 7.46-7.52 (m, 2$H_{arom}$), 7.32-7.43 (m, 5$H_{arom}$), 7.22-7.26 (m, 1$H_{arom}$), 7.13-7.18 (m, 1$H_{arom}$), 3.90 (t, *J*=8Hz, 1H), 3.29 (s, 2H), 3.27 (s, 6H), 2.49 (d, *J*=8Hz, 2H). $^{13}$C NMR ($CDCl_3$): 169.6 (s, 1C=O), 138.6 (s, 1q$C_{arom}$), 134.8 (s, 1q$C_{arom}$), 132.0 (s, 1q$C_{arom}$), 130.1 (s, 1$C_{arom}$), 129.4 (s, 2$C_{arom}$), 128.9 (s, 2$C_{arom}$), 128.5 (s, 1$C_{arom}$), 127.7 (s, 1$C_{arom}$), 123.9 (s, 1$C_{arom}$), 120.4 (s, 1$C_{arom}$), 103.1 (s, 1CH), 53.9 (s, 2$CH_3$), 52.8 (s, 1$CH_2$), 50.9 (s, 1$CH_2$). MS (EI): m/z (%): 315 (1) $[M+H]^+$, 282 (77), 239 (33), 182 (45), 167 (20), 152 (5), 139 (3), 118 (49), 86 (100), 75 (85), 58 (15), 42 (22).

**1,2,3,13b-tetrahydropyrazino[1,2-f]phenanthridin-4-one**

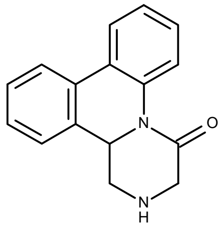


A solution of 5g 2-(2,2-dimethoxyethylamino)-*N*-(2-phenylphenyl)acetamide (15.9mmol) in DCM (15mL) is slowly added to a mixture of 5.6mL trifluoro-methane sulfonic acid (63.6mmol) in DCM (25mL) cooled down at 0°C under nitrogen. The mixture is then stirred overnight at RT before adding water (40mL). The phases are separated, the aqueous phase is neutralized with 32% aqueous sodium hydroxide solution until the pH reaches 10-11 and is subsequently extracted 3 times with DCM (40mL). The combined organic phases are washed with water (100mL) and toluene (100mL) is added before evaporation under vacuum. The evaporation residue is purified by column chromatography (EtOAc/methanol 9/1 and 7/3) affording the target compound as a slightly brown solid (0.68g, 2.7mmol, 17% yield).

$C_{16}H_{14}N_2O$ (250.29g.mol$^{-1}$) m.p. 148-149°C. $^1$H NMR ($CDCl_3$): 7.79-7.86 (m, 3$H_{arom}$), 7.42-7.47 (m, 1$H_{arom}$), 7.35-7.41 (m, 2$H_{arom}$), 7.29-7.33 (m, 2$H_{arom}$), 4.60 (t, *J*=5Hz, 1H), 3.81 (d, *J*=20Hz, 1H), 3.69-3.80 (m, 2H), 3.54 (d, *J*=20Hz, 1H). $^{13}$C NMR ($CDCl_3$): 167.2 (s, 1C=O), 136.7 (s, 1q$C_{arom}$), 134.9 (s, 1q$C_{arom}$), 133.0 (s, 1q$C_{arom}$), 128.5 (s, 1q$C_{arom}$), 128.4 (s, 1$C_{arom}$), 128.1 (s, 1$C_{arom}$), 128.0 (s, 1$C_{arom}$), 126.1 (s, 1$C_{arom}$), 125.3 (s, 1$C_{arom}$), 124.3 (s, 1$C_{arom}$), 124.1 (s, 1$C_{arom}$), 122.2 (s, 1$C_{arom}$), 56.0 (s, 1CH), 51.2 (s, 1$CH_2$), 44.6 (s, 1$CH_2$)

MS (EI): m/z (%): 250 (20) [M]$^+$, 221 (41), 193 (14), 180 (100), 165 (5), 152 (18), 139 (2), 126 (3), 95 (4), 84 (6), 76 (4), 63 (4), 49 (6), 43 (35), 28 (5).

**2-(cyclohexanecarbonyl)-3,13b-dihydro-1*H*-pyrazino[1,2-f]phenanthridin-4-one (34)**

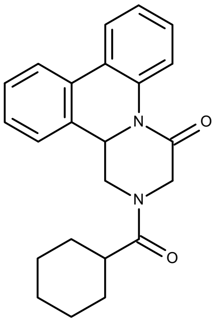


1.26g 1,2,3,13b-tetrahydropyrazino[1,2-f]phenanthridin-4-one (5.03mmol) are dissolved in a biphasic mixture of DCM (30mL) and water (30mL) at RT under nitrogen. The pH is set to 10 by adding a few drops of 32% aqueous sodium hydroxide solution. A solution of 0.74g cyclohexane carboxylic acid chloride (5.03mmol) in DCM (10mL) and 0.49mL of 32% aqueous sodium hydroxide solution are then slowly added in parallel keeping the pH between 9 and 11. After addition, the mixture is further stirred for 2h before phase separation. The aqueous phase is extracted 3 times with DCM (20mL), the combined organic phases are washed 3 times with a 5% aqueous ammonia solution and finally with water until the pH reaches 9. After evaporation of the solvent, the residue is purified by column chromatography (EtOAc/*n*-heptane 1/1) affording the target compound as a white solid (0.56g, 1.55mmol, 31% yield).

$C_{23}H_{24}N_2O_2$ (360.45g.mol$^{-1}$) m.p. 134-139°C. $^1$H NMR ($CDCl_3$): 7.77-7.86 (m, 3$H_{arom}$), 7.48-7.53 (m, 1$H_{arom}$), 7.33-7.46 (m, 4$H_{arom}$), 5.07 (dd, *J*=16, 4Hz, 1H), 4.63-4.68 (m, 1H), 4.27-4.38 (m, 2H), 3.92 (dd, *J*=16, 4Hz, 1H), 2.44 (t, *J*=12Hz, 1H), 1.65-1.86 (m, 4H), 1.41-1.61 (m, 2H), 1.17-1.32 (m, 4H). $^{13}$C NMR ($CDCl_3$): 174.7 (s, 1C=O), 163.6 (s, 1C=O), 136.4 (s, 1q$C_{arom}$), 133.9 (s, 1q$C_{arom}$), 132.4 (s, 1q$C_{arom}$), 128.8 (s, 1q$C_{arom}$), 128.6 (s, 1$C_{arom}$), 128.4 (s, 1$C_{arom}$), 128.0 (s, 1$C_{arom}$), 126.7 (s, 1$C_{arom}$), 125.1 (s, 1$C_{arom}$), 124.3 (s, 1$C_{arom}$), 124.1 (s, 1$C_{arom}$), 123.0 (s, 1$C_{arom}$), 56.0 (s, 1CH), 49.7 (s, 1$CH_2$), 40.6 (s, 2$CH_2$), 38.7 (s, 1CH), 29.3 (s, 1$CH_2$), 28.8 (s, 1$CH_2$), 25.6 (s, 2$CH_2$). MS (EI): m/z (%): 360 (14) [M]$^+$, 249 (5), 221 (8), 193 (27), 180 (100), 165 (8), 152 (23), 140 (5), 126 (38), 113 (75), 98 (6), 83 (36), 67 (5), 55 (46), 42 (30), 32 (10).

**4b,6,10b,12-Tetrahydro-5a,11a-diaza-indeno[1,2-b]fluorene-5,11-dione (43)**

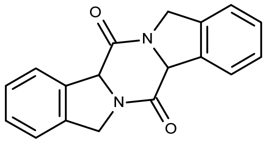


$C_{18}H_{14}N_2O_2$ (290.32 g.mol$^{-1}$), m.p. 219-222°C. $^1$H NMR ($CDCl_3$): 7.65-7.69 (m, 2$H_{arom}$), 7.38-7.43 (m, 6$H_{arom}$), 5.76 (s, 2H), 4.78 (d, *J*=16Hz, 2H), 4.63 (d, *J*=16Hz, 2H). $^{13}$C NMR ($CDCl_3$): 165.1 (s, 2C=O), 136.6 (s, 2q$C_{arom}$), 134.4 (s, 2q$C_{arom}$), 128.7 (s, 2$C_{arom}$), 127.7 (s, 2$C_{arom}$), 125.9 (s, 2$C_{arom}$), 123.4 (s, 2$C_{arom}$), 63.5 (s, 2CH), 50.4 (s, 2$CH_2$)

**2-(4-Butyl-cyclohexanecarbonyl)-1,2,3,6,7,11b-hexahydro-pyrazino[2,1-a]isoquinolin-4-one (11)**

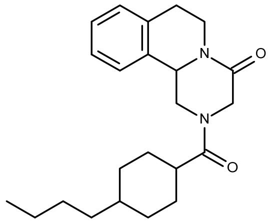

$C_{23}H_{32}N_2O_2$ (368.51 g.mol$^{-1}$), m.p. 139-140°C. $^{1}$H NMR ($CDCl_3$): 7.20-7.31 (m, 4$H_{arom}$), 4.77-4.96 (m, 1H), 4.38-4.56 (m, 3H), 3.73 (d, J=16Hz, 1H), 3.29 (d, *J*=16Hz, 2H), 2.75-2.90 (m, 4H), 1.63-1.78 (m, 4H), 1.16-1.43 (m, 8H), 0.96-1.11 (m, 2H), 0.87 (t, *J*=8Hz, 3H). $^{13}$C NMR ($CDCl_3$): 174.25 (s, 1C=O), 164.9 (s, 1C=O), 135.5 (s, 1q$C_{arom}$), 133.8 (s, 1q$C_{arom}$), 129.4 (s, 1$C_{arom}$), 127.5 (s, 1$C_{arom}$), 127.0 (s, 1$C_{arom}$), 126.2 (s, 1$C_{arom}$), 54.9 (s, 1CH), 48.7 (s, 1$CH_2$), 45.6 (s, 1$CH_2$), 39.0 (s, 1CH), 37.0 (s, 2$CH_2$), 32.1 (s, 1$CH_2$), 29.6 (s, 1CH), 29.2 (s, 1$CH_2$), 29.0 (s, 2$CH_2$), 28.6 (s, 1$CH_2$), 22.9 (s, 2$CH_2$), 14.5 (s, 1$CH_3$).

**Fig. S1.**

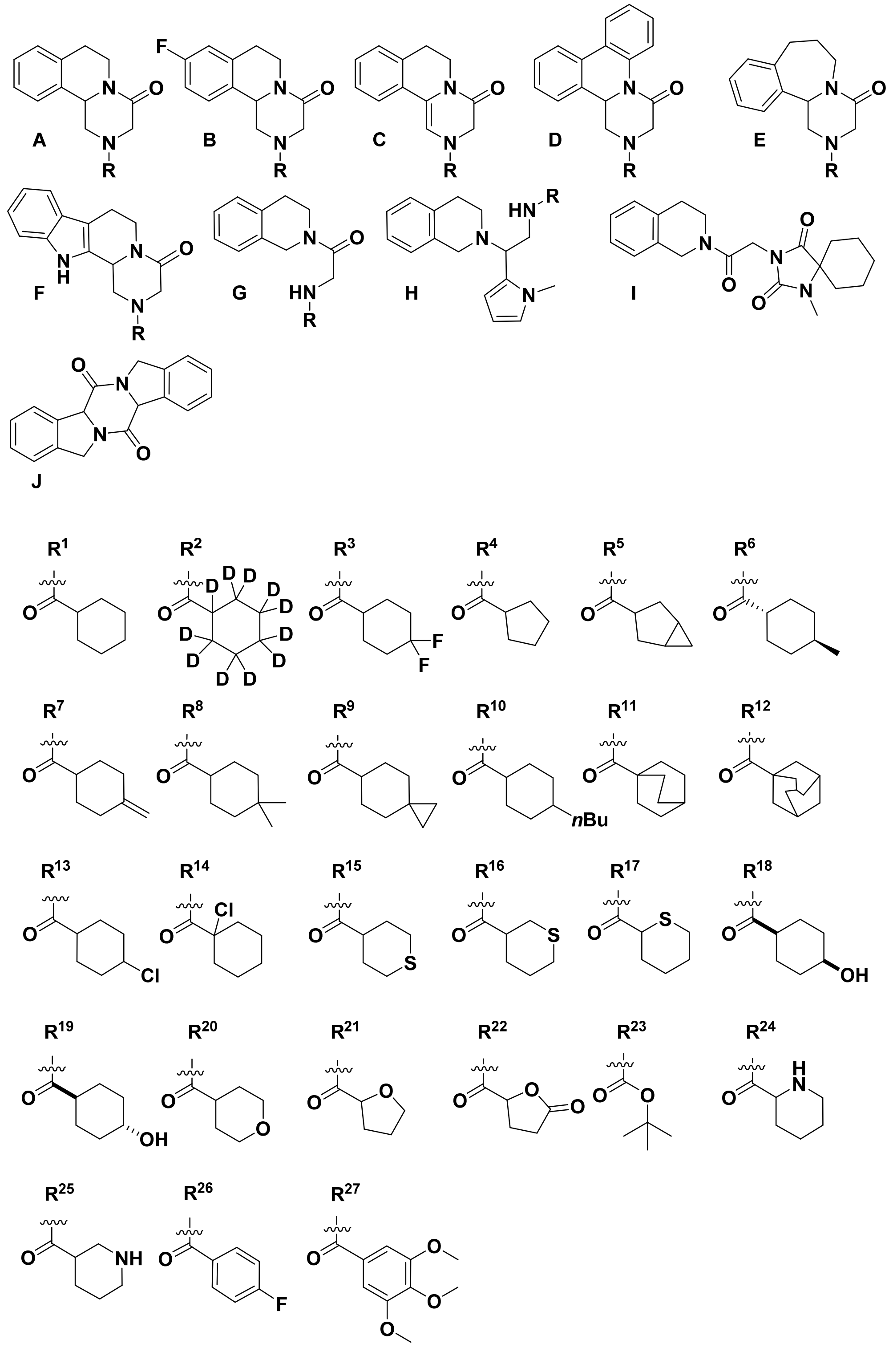


**Fig. S1. Structures of analogues**. The series of 43 analogues used in this study represented combinations of core (A through J, top) and substituent groupings ($R^1$ through $R^{27}$, bottom) as detailed in Table S1. Analogues are referred to by number (Table S1) and composition to include stereochemistry. For example, (*R*)-PZQ is represented by [A(*R*)$R^1$] and (*S*)-PZQ is represented by [A(*S*)$R^1$].

**Fig. S2.**

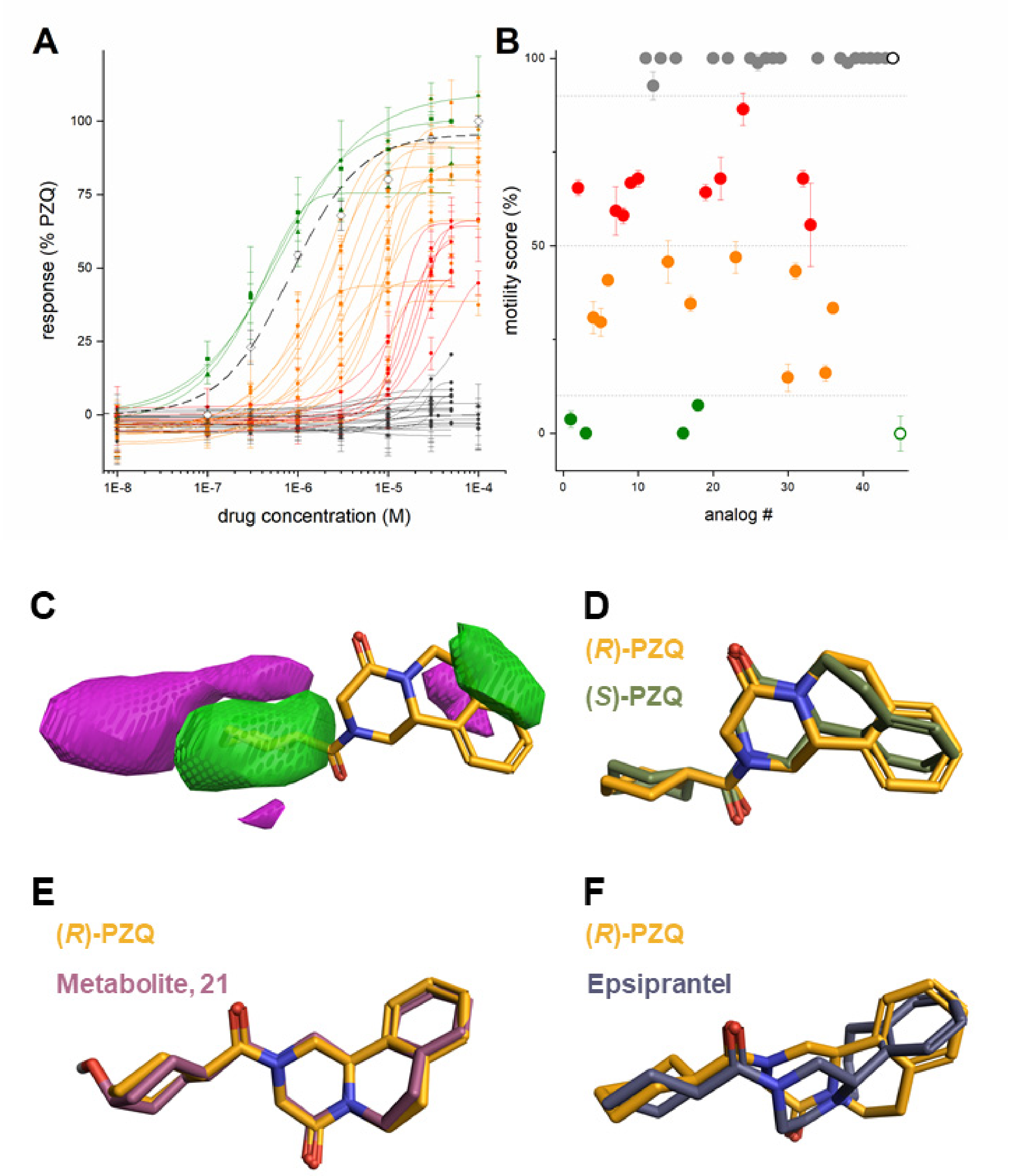


**Fig. S2**. **Ligand-based activity model derived from target and phenotypic analogue analyses.** (**A**) Dose response relationships for a series of 43 analogues (average $M_r$=330, average cLogP=2.24) profiled against *Sm*.$TRPM_{PZQ}$ in a fluorometric $Ca^{2+}$ assay. Curves are illustrated in different colors to illustrate analogues with different potency from high (green) to 'inactive' (grey) as defined in the main text. For clarity, the identity of each analogue (Table S1) is not indexed except for ±PZQ (dashed black line). Results represent the average of technical replicates from at least three independent transfections and are normalized to the peak fluorescence value evoked by ±PZQ (100µM). (**B**) Effect of analogues on schistosome worm contraction. Quantification of movement assays for each analogue. Data are expressed relative to control (no drug, mobility score=100, open black circle) and PZQ-treatment (mobility score=0, open green circle). Data are averaged across tested concentrations and timepoints and the mean mobility score categorized by color: green (<10% of control value), orange, (10-50% of control value), red (50-90% of control value) or lack of activity (grey). (**C**) Summary of favorable (green) and unfavorable (purple) hydrophobic interactions obtained from SAR data. The cyclohexyl ring is associated with favorable hydrophobic interactions, whereas the area surrounding it is associated with unfavorable interactions. The pyrazinoisoquinoline core is covered by small areas of favorable (front) and unfavorable interactions (behind). (**D**) Structural alignment of (*R*)-PZQ (orange) and (*S*)-PZQ (green). When the cyclohexyl rings of both enantiomers are well aligned, the pyrazinoisoquinoline core of (*S*)-PZQ is twisted out of the plane the (*R*)-PZQ core. (**E**) Alignment between (*R*)-PZQ (orange) and the metabolite analog 21 (purple). Both structures overlap almost perfectly, only the hydroxyl substituent at the cyclohexyl ring is the differential feature. (**F**) Overlap of (*R*)-PZQ (orange) and epsiprantel (blue). Epsiprantel core structure is shifted compared to pyrazinoisoquinoline core of (*R*)-PZQ.

**Fig. S3.**

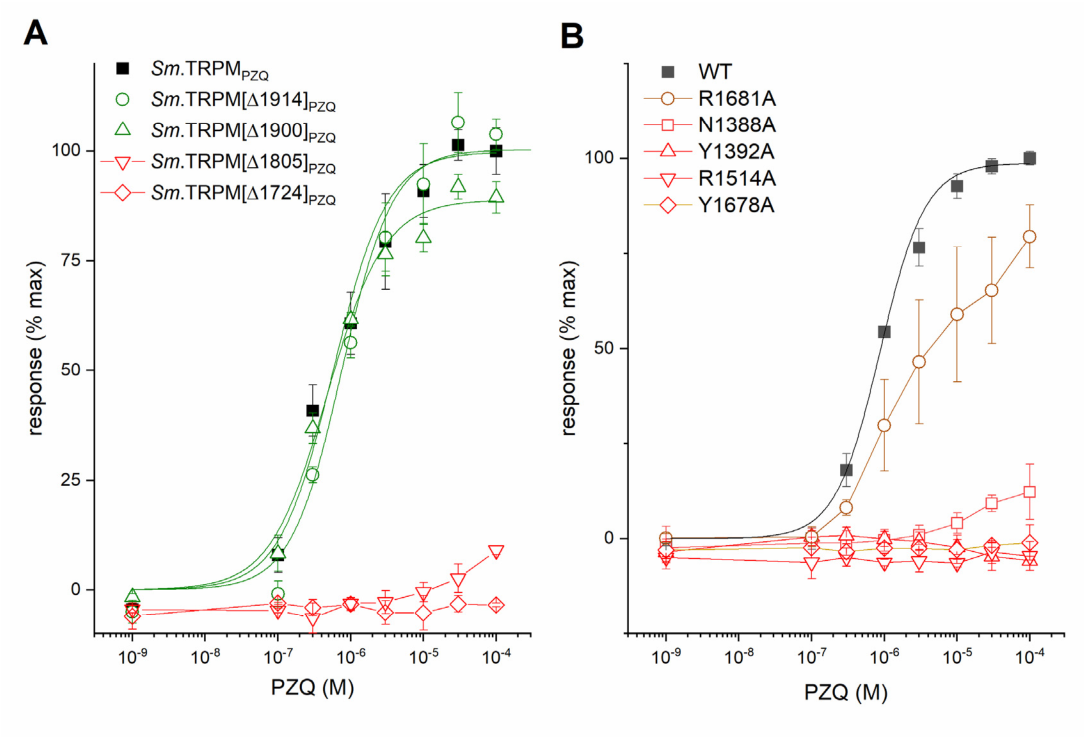


**Fig. S3. Mutagenesis of *Sm*.TRPM$_{PZQ}$.** (**A**) Concentration-response relationship for wild-type *Sm*.TRPM$_{PZQ}$ (solid square) and COOH terminal truncation mutants (open symbols) to ±PZQ. (**B**) Concentration-response relationship for wild-type *Sm*.TRPM$_{PZQ}$ (solid square) and indicated point mutants in response to ±PZQ. Data represent mean±sem from an average of three independent transfections.

**Fig. S4.**

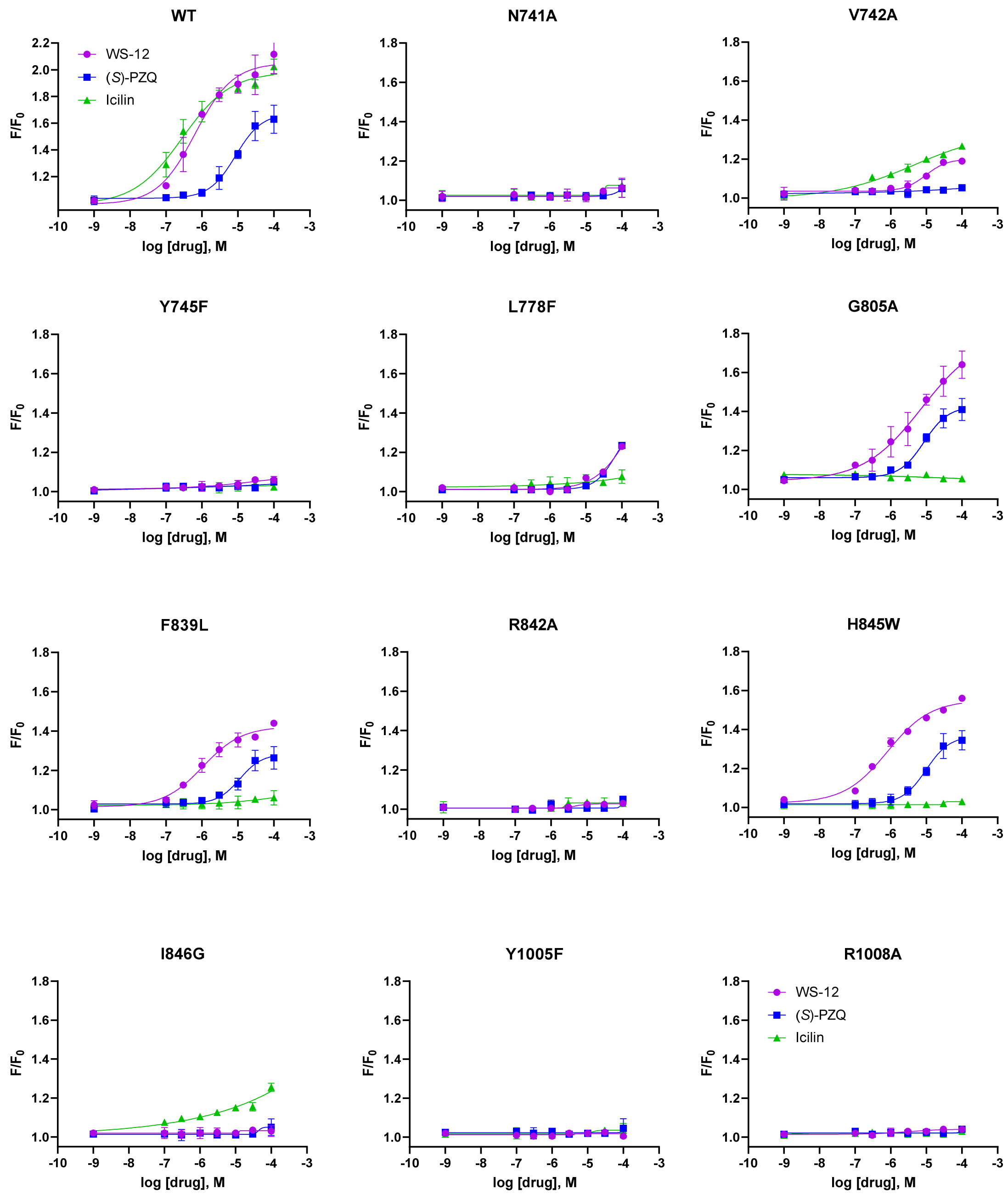


**Fig. S4. Mutational analysis of binding pocket residues in human TRPM8.** Concentration-response curves of wild type human hTRPM8, and 11 different hTRPM8 point mutants in response to (*S*)-PZQ (blue square), icilin (green triangle) and WS-12 (purple circle). Responses were measured in intact cells using the $Ca^{2+}$-sensitive fluorescent dye fluo-4. Data represent mean±sem from an average of three independent transfections.

**Fig. S5.**

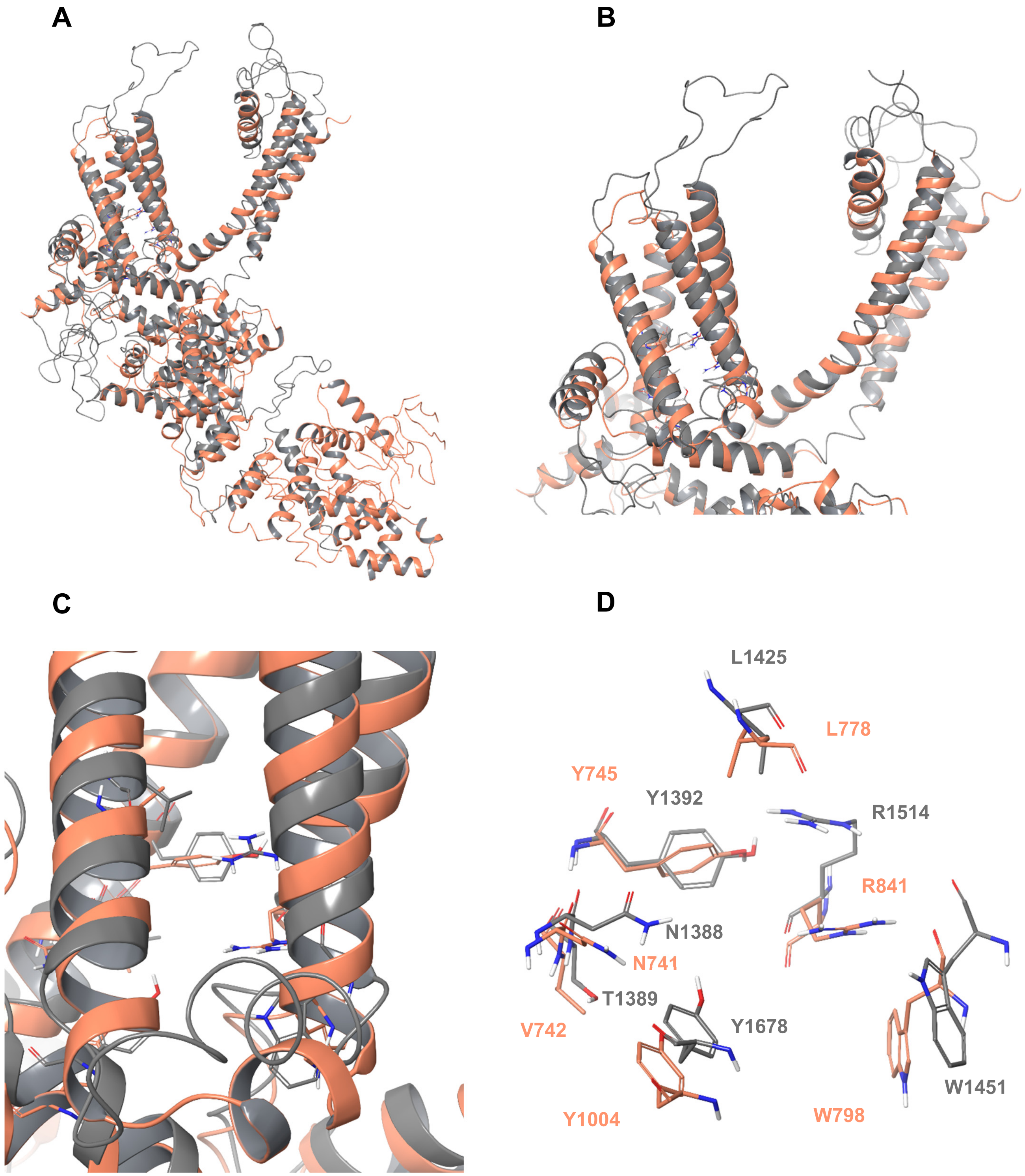


**Fig. S5. Comparison of *Sm*.TRPM$_{PZQ}$ homology model with human TRPM8 structure**. (**A**) Overall structure superposition of *Sm*.TRPM$_{PZQ}$ homology model (grey) with the human TRPM8 structure (PDB ID: 6NR3, orange) based on sequence alignment (root mean square difference (RMSD) of all atoms is 9.06Å). (**B**) Enlargement of transmembrane region to show superimposed VSLD (left) and pore domain (right). (**C**) VSLD (transmembrane helices S1-S4) and known TRPM8 ligand-binding residues superimposed with the aligned residues from the *Sm*.$TRPM_{PZQ}$ model. (**D**) Annotated residues known for ligand interactions of TRPM8 activators and the equivalent superimposed *Sm*.$TRPM_{PZQ}$ residues.

**Fig. S6.**

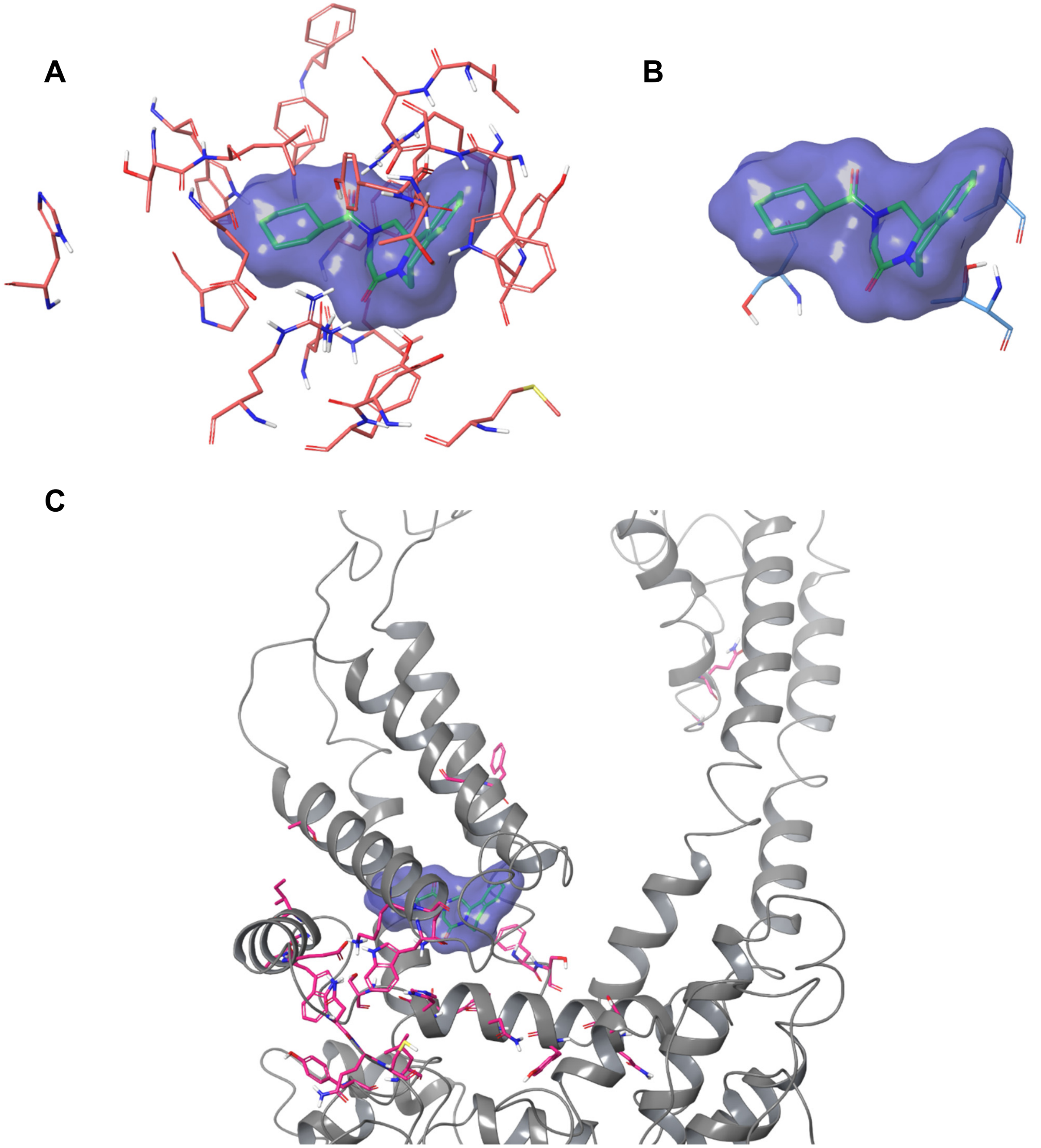


**Fig. S6. Location of residues mutated in functional analysis of *Sm*.TRPM$_{\mathbf{PZQ}}$.** (**A**) Twenty residues that lead to decreased PZQ sensitivity when mutated to alanine (light red) lie close to the predicted PZQ binding site. The docked PZQ structure is highlighted in green and its ligand surface is shown in blue. (**B**) Location of three residues that do not affect PZQ sensitivity when mutated to alanine (S1391A, L1515A and T1518A, light blue). (**C**), Location of other residues (purple) targeted for mutation outside the predicted binding pocket area in the homology model.

**Table S1.**

| | | *Core* | *Stereo-* | *R* | *$EC_{50}$ (µM)* | cLogP | *$M_r$* | *Reference* |
|---|---|---|---|---|---|---|---|---|
| *1* | (*R*)-PZQ | A | *R* | $R^1$ | 0.5 ± 0.1 | 2.407 | 312.4 | (*9*) |
| *2* | (*S*)-PZQ | A | *S* | $R^1$ | 24.7 ± 1.3 | 2.407 | 312.4 | (*9*) |
| *3* | | A | *R* | $R^2$ | 0.3 ± 0.02 | 2.407 | 323.4 | (*33*) |
| *4* | | A | *R* | $R^3$ | 2.5 ± 0.7 | 2.284 | 348.3 | (*34*) |
| *5* | | A | *R* | $R^4$ | 3.2 ± 1.8 | 2.172 | 298.3 | SM |
| *6* | | A | *R* | $R^5$ | 3.2 ± 1.1 | 2.383 | 310.3 | SM |
| *7* | | A | *R* | $R^6$ | 9.6 ± 0.8 | 2.952 | 326.4 | SM |
| *8* | | A | *R* | $R^7$ | 7.5 ± 1.1 | 2.455 | 324.4 | SM |
| *9* | | A | *R* | $R^8$ | 17.0 ± 3.1 | 3.175 | 340.4 | SM |
| *10* | | A | *R* | $R^9$ | 19.5 ± 3.0 | 2.709 | 338.4 | SM |
| *11* | | A | racemic | $R^{10}$ | inactive | 3.564 | 368.5 | SM |
| *12* | | A | *R* | $R^{11}$ | inactive | 2.845 | 338.4 | SM |
| *13* | | A | racemic | $R^{12}$ | inactive | 2.982 | 364.4 | (*34*) |
| *14* | | A | *R* | $R^{13}$ | 5.3 ± 1.8 | 2.670 | 346.8 | SM |
| *15* | | A | *R* | $R^{14}$ | inactive | 2.832 | 346.8 | SM |
| *16* | | A | *R* | $R^{15}$ | 0.3 ± 0.01 | 2.100 | 330.4 | SM |
| *17* | | A | *R* | $R^{16}$ | 2.5 ± 0.6 | 2.100 | 330.4 | SM |
| *18* | | A | *R* | $R^{17}$ | 1.6 ± 0.5 | 2.100 | 330.4 | SM |
| *19* | *cis*-hydroxy-(*R*)-PZQ | A | *R* | $R^{18}$ | 13.4 ± 2.9 | 1.327 | 328.4 | (*9*) |
| *20* | *cis*-hydroxy-(*S*)-PZQ | A | *S* | $R^{18}$ | inactive | 1.327 | 328.4 | (*9*) |
| *21* | *trans*-hydroxy-(*R*)-PZQ | A | *R* | $R^{19}$ | 18.0 ± 2.0 | 1.327 | 328.4 | (*9*) |
| *22* | *trans*-hydroxy-(*S*)-PZQ | A | *S* | $R^{19}$ | inactive | 1.327 | 328.4 | (*9*) |
| *23* | | A | *R* | $R^{20}$ | 8.1 ± 0.7 | 0.964 | 314.3 | (*34*) |
| *24* | | A | *R* | $R^{21}$ | 33.3 ± 6.4 | 0.736 | 300.3 | SM |
| *25* | | A | *R* | $R^{22}$ | inactive | 0.468 | 314.3 | SM |
| *26* | | A | *R* | $R^{23}$ | inactive | 2.667 | 302.3 | SM |
| *27* | | A | *S* | $R^{23}$ | inactive | 2.667 | 302.3 | SM |
| *28* | | A | *R* | $R^{24}$ | inactive | 1.301 | 313.3 | SM |
| *29* | | A | *R* | $R^{25}$ | inactive | 0.903 | 313.3 | SM |
| *30* | | B | *R* | $R^1$ | 1.3 ± 0.2 | 2.526 | 330.4 | SM |
| *31* | | B | *R* | $R^4$ | 4.0 ± 1.1 | 2.236 | 316.3 | SM |
| *32* | | B | racemic | $R^{20}$ | 13.6 ± 0.4 | 1.007 | 332.3 | SM |
| *33* | | C | NA | $R^1$ | 13.7 ± 1.8 | 2.037 | 310.3 | (*35*) |
| *34* | | D | racemic | $R^1$ | inactive | 3.123 | 360.4 | SM |
| *35* | epsiprantel | E | racemic | $R^1$ | 1.6 ± 0.1 | 2.643 | 326.4 | (*36*) |
| *36* | | E | racemic | $R^4$ | 4.0 ± 0.4 | 2.355 | 312.4 | (*36*) |
| *37* | | F | racemic | $R^1$ | inactive | 2.948 | 351.4 | (*37*) |
| *38* | | F | racemic | $R^{26}$ | inactive | 2.991 | 363.3 | (*37*) |
| *39* | | F | racemic | $R^{27}$ | inactive | 2.699 | 435.4 | (*37*) |
| *40* | | G | NA | $R^1$ | inactive | 1.718 | 300.4 | CAS879730-20-88 |
| *41* | | H | NA | $R^1$ | inactive | 3.904 | 365.5 | CAS1049459-55-3 |
| *42* | | I | NA | - | inactive | 2.416 | 355.4 | CAS1043202-32-9 |
| *43* | | J | NA | NA | inactive | 2.198 | 290.3 | SM |

**Table S1. Overview of analogues used in *Sm*.TRPM$_{PZQ}$ assays and worm experiments.** List of 43 analogues ordered by core structure ('A' through J') and substituents ($R^1$ through $R^{27}$) as shown in Figure S1. Key properties (stereochemistry, calculated LogP (cLogP) and molecular weight ($M_r$)) of each analogue are defined. $EC_{50}$s for activation of *Sm*.TRPM$_{PZQ}$ are listed, and analogues displaying either no measurable activity or activity ≤25% of PZQ at 50µM are categorized as 'inactive'. Synthetic procedures for each analogue are either indexed, referenced or detailed in the Supplementary Material and Methods ('SM').

**Table S2.**

| | Residue | Mutant | Location | $EC_{50}$ (µM) | $B_{max}$ (% WT) |
|---|---|---|---|---|---|
| 1 | T109 | T109A | MHR | 0.76 ± 0.13 | 125.9 ± 8.2 |
| 2 | D110 | D110A | MHR | 0.89 ± 0.21 | 127.9 ± 5.6 |
| 3 | E112 | E112A | MHR | 0.28 ± 0.04 | 115.2 ± 3.1 |
| 4 | M145 | M145A | MHR | 0.58 ± 0.1 | 67.1 ± 8.8 |
| 5 | G216 | G216A | MHR | 1.21 ± 0.71 | 89.3 ± 19.3 |
| 6 | I218 | I218A | MHR | 0.53 ± 0.1 | 81.1 ± 4.2 |
| 7 | K219 | K219A | MHR | 1.09 ± 0.3 | 98.1 ± 14.0 |
| 8 | R222 | R222A | MHR | 0.59 ± 0.01 | 76.0 ± 8.4 |
| 9 | D224 | D224A | MHR | 0.78 ± 0.12 | 110.4 ± 2.8 |
| 10 | E226 | E226A | MHR | 0.84 ± 0.23 | 95.3 ± 6.2 |
| 11 | R228 | R228A | MHR | 0.62 ± 0.1 | 105.3 ± 7.6 |
| 12 | T244 | T244A | MHR | 0.76 ± 0.14 | 118.0 ± 9.1 |
| 13 | T245 | T245A | MHR | 0.90 ± 0.17 | 179.7 ± 20.1 |
| 14 | D246 | D246A | MHR | 0.53 ± 0.21 | 78.0 ± 6.0 |
| 15 | E252 | E252A | MHR | 0.78 ± 0.2 | 175.8 ± 6.0 |
| 16 | E254 | E254A | MHR | 0.73 ± 0.1 | 81.0 ± 3.1 |
| 17 | R258 | R258A | MHR | 0.62 ± 0.13 | 138.0 ± 10.2 |
| 18 | F992 | F992L | MHR | 0.49 ± 0.1 | 74.9 ± 1.0 |
| 19 | K1205 | K1205Q | MHR | 0.69 ± 0.1 | 73.2 ± 11.5 |

**Table S2. Mutational analysis of residues within the MHR domain of *Sm*.TRPM$_{PZQ}$.**
PZQ sensitivity of individual point mutants localized with the $NH_2$-terminal MHR domains of *Sm*.TRPM$_{PZQ}$. For ease of comparison between supplementary tables, responsiveness of each mutant is categorized by color, to reflect mutants exhibiting an $EC_{50}$ ≤1µM (green), 1-3µM (orange) and ≥3µM or irresponsive (red). $B_{max}$ represents peak response relative to response evoked by PZQ (100µM) on wild type *Sm*.TRPM$_{PZQ}$.

**Table S3.**

| | Residue | Mutant | Location | $EC_{50}$ (µM) | $B_{max}$ (% WT) |
|---|---|---|---|---|---|
| **1** | N1388 | N1388A | S1 | No signal | 7.2 ± 3.5 |
| **2** | T1389 | T1389A | S1 | 1.85 ± 0.84 | 46.6 ± 3.1 |
| **3** | S1391 | S1391A | S1 | 0.83 ± 0.28 | 75.1 ± 4.2 |
| **4** | Y1392 | Y1392A | S1 | No signal | 4.8 ± 2.8 |
| **5** | F1395 | F1395A | S1 | No signal | 1.2 ± 1.2 |
| **6** | W1421 | W1421A | S2 | No signal | 8.4 ± 3.1 |
| **7** | T1424 | T1424A | S2 | 1.39 ± 0.29 | 82.1 ± 5.3 |
| **8** | L1425 | L1425A | S2 | 2.41 ± 0.41 | 61.9 ± 9.0 |
| | | L1425F | S2 | 6.0±3.8 | 45.3±8.0 |
| | | L1425W | S2 | 30.4±5.1 | 7.6±5.5 |
| **9** | E1428 | E1428A | S2 | No signal | 2.1 ± 0.9 |
| **10** | W1451 | W1451A | S3 | No signal | -4.7 ± 2.9 |
| **11** | N1452 | N1452A | S3 | 2.5 ± 0.9 | 41.9 ± 2.8 |
| **12** | L1454 | L1454A | S3 | 1.8 ± 0.38 | 99.2 ± 4.8 |
| **13** | D1455 | D1455A | S3 | No signal | 3.7 ± 0.3 |
| **14** | F1511 | F1511A | S4 | No signal | -7.4 ± 2.5 |
| **15** | R1514 | R1514A | S4 | No signal | -9.5 ± 2.2 |
| **16** | L1515 | L1515A | S4 | 0.7 ± 0.03 | 86.2 ± 7.9 |
| **17** | Y1517 | Y1517A | S4 | No signal | 1.1 ± 2.3 |
| **18** | T1518 | T1518A | S4 | 0.53 ± 0.2 | 121.8 ± 11.1 |
| **19** | D1677 | D1677A | TRP | 1.4 ± 0.3 | 27.0 ± 1.0 |
| **20** | Y1678 | Y1678A | TRP | No signal | -3.1 ± 0.2 |
| **21** | R1681 | R1681A | TRP | 2.1 ± 0.7 | 63.0 ± 7.1 |
| **22** | P1686 | P1686A | TRP | 6.7 ± 2.2 | 18.2 ± 1.1 |
| **23** | I1689 | I1689A | TRP | 3.6 ± 0.8 | 109.3 ± 31.3 |

**Table S3. Mutational analysis of *Sm*.TRPM$_{PZQ}$ residues lining the predicted PZQ binding pocket**. PZQ sensitivity of individual point mutants that line the predicted PZQ binding pocket measured in a fluorimetric $Ca^{2+}$ assay. Responsiveness of each mutant is categorized by color, to reflect mutants exhibiting an $EC_{50}$ ≤1µM (green), 1-3µM (orange) and ≥3µM, or irresponsive (red).

**Table S4.**

| | Residue | Mutant | Location | $EC_{50}$ (µM) | $B_{max}$ (% WT) | Distance from binding pocket (Å) |
|---|---|---|---|---|---|---|
| 1 | Y1347 | Y1347A | pre-S1 | 0.97 ± 0.09 | 108.2 ± 14.1 | 21.63 |
| 2 | L1349 | L1349A | pre-S1 | 1.02 ± 0.23 | 99.97 ± 10.0 | 17.78 |
| 3 | F1385 | F1385A | S1 | 0.72 ± 0.09 | 85.6 ± 10.3 | 9.11 |
| 4 | A1420 | A1420L | S2 | 0.89 ± 0.4 | 135.3 ± 4.7 | 16.46 |
| 5 | K1431 | K1431A | S2 | 1.78 ± 0.3 | 77.2 ± 3.4 | 13.66 |
| 6 | Q1432 | Q1432A | S2 | 1.03 ± 0.37 | 70.4 ± 4.0 | 11.98 |
| 7 | W1435 | W1435A | S2 | 0.9 ± 0.2 | 155.1 ± 5.1 | 16.76 |
| 8 | G1458 | G1458A | S3 | 1.2 ± 0.4 | 98.1 ± 9.4 | 12.22 |
| 9 | F1509 | F1509A | S4 | 1.4 ± 0.33 | 76.9 ± 5.8 | 14.30 |
| 10 | F1519 | F1519A | S4 | 2.6 ± 0.8 | 103.3 ± 12.0 | 9.64 |
| 11 | S1520 | S1520A | S4-S5 | 2.9 ± 0.64 | 51.6 ± 2.1 | 10.56 |
| 12 | Q1598 | Q1598A | S5-S6 | 0.55 ± 0.2 | 99.2 ± 1.3 | 38.42 |
| 13 | G1601 | G1601A | S5-S6 | 0.65 ± 0.1 | 117.6 ± 0.7 | 37.87 |
| 14 | N1668 | N1668A | TRP | 0.64 ± 0.2 | 101.1 ± 2.7 | 19.63 |
| 15 | Y1669 | Y1669A | TRP | 0.73 ± 0.1 | 64.5 ± 5.7 | 19.17 |
| 16 | Y1672 | Y1672A | TRP | 1.0 ± 0.7 | 42.2 ± 5.7 | 16.54 |
| 17 | N1676 | N1676A | TRP | 0.46 ± 0.1 | 121.1 ± 2.6 | 13.96 |
| 18 | H1680 | H1680A | TRP | 1.60 ± 0.3 | 96.6 ± 5.4 | 13.56 |
| 19 | S1682 | S1682A | TRP | 0.55 ± 0.2 | 85.6 ± 5.1 | 13.24 |
| 20 | I1688 | I1688A | TRP | 0.8 ± 0.2 | 129.8 ± 20.9 | 15.20 |
| 21 | I1690 | I1690A | TRP | 1.1 ± 0.1 | 83.7 ± 7.1 | 15.87 |
| 22 | W1692 | W1692A | TRP | 0.76 ± 0.09 | 87.6 ± 4.7 | 17.39 |
| 23 | E1696 | E1696A | TRP | 1.2 ± 0.24 | 85.3 ± 6.6 | 20.41 |
| 24 | N1703 | N1703A | COOH | 1.0 ± 0.2 | 129.2 ± 10.9 | 24.70 |
| 25 | Q1704 | Q1704A | COOH | 1.5 ± 0.5 | 114.1 ± 6.6 | 24.70 |
| 26 | C1705 | C1705A | COOH | 1.5 ± 0.4 | 79.0 ± 6.5 | 22.82 |

**Table S4. Mutational analysis of *Sm*.TRPM$_{PZQ}$ residues localized outside predicted PZQ binding pocket**. PZQ sensitivity of individual point mutants in regions of *Sm*.TRPM$_{PZQ}$ located outside the predicted PZQ binding pocket. Responsiveness of each mutant is categorized by color, to reflect mutants exhibiting an $EC_{50}$ ≤1µM (green), 1-3µM (orange) and ≥3µM, or irresponsive (red). Distance is calculated between residues and centroid of piperazin-2-one ring of PZQ.

**Table S5.**

| | 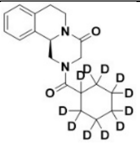 | 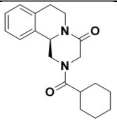 | 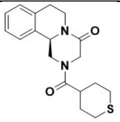 | 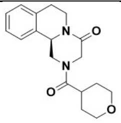 | 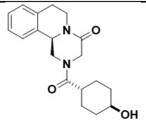 |
|---|---|---|---|---|---|
| **Analog ID** | **3** | **1** | **16** | **23** | **21** |
| **LogP** | 2.407 | 2.407 | 2.100 | 0.964 | 1.327 |
| **TPSA (Å$^2$)** | 40.62 | 40.62 | 65.92 | 49.85 | 60.85 |
| ***Sm.*TRPM$_{PZQ}$ IC$_{50}$ (µM)** | 0.3 ± 0.02 | 0.5 ± 0.1 | 0.3 ± 0.01 | 8.1 ± 0.7 | 18.0 ± 2.0 |
| **Human microsomal CL$_{int}$ (µL/min/mg prot)** | 34 | 63 | 28 | <10 | ND |
| **Mouse microsomal CL$_{int}$ (µL/min/mg prot)** | 818 | >1000 | 324 | 17 | ND |

**Table S5**. **Metabolic profile of PZQ analogs**. Metabolic profile of indicated analogues (4, 1, 16, 20 & 21) measured *in vitro* using human and mouse microsomes. For each of the indicated compounds, *in vitro* intrinsic clearance (Cl$_{int}$) was analyzed using human and mouse liver microsome enzymes. EC$_{50}$ values at *Sm*.TRPM$_{PZQ}$, calculated LogP and TPSA are also shown. ND, not determined.

**Table S6.**

| | | *Core* | *Stereo-* | *R* | *EC50 (µM)* |
|---|---|---|---|---|---|
| *1* | (*R*)-PZQ | A | *R* | $R^1$ | 0.15±0.05 |
| *2* | (*S*)-PZQ | A | *S* | $R^1$ | 9.87±0.75 |
| *3* | | A | *R* | $R^2$ | 0.11±0.06 |
| *4* | | A | *R* | $R^3$ | 1.4±0.46 |
| *5* | | A | *R* | $R^4$ | 0.63±0.24 |
| *6* | | A | *R* | $R^5$ | 1.33±0.61 |
| *7* | | A | *R* | $R^6$ | 4.8±0.95 |
| *8* | | A | *R* | $R^7$ | 1.33±0.29 |
| *9* | | A | *R* | $R^8$ | 10.10±0.1 |
| *10* | | A | *R* | $R^9$ | 9.35±0.5 |
| *11* | | A | racemic | $R^{10}$ | inactive |
| *12* | | A | *R* | $R^{11}$ | inactive |
| *13* | | A | racemic | $R^{12}$ | inactive |
| *14* | | A | *R* | $R^{13}$ | 1.85±0.53 |
| *15* | | A | *R* | $R^{14}$ | inactive |
| *16* | | A | *R* | $R^{15}$ | 0.08±0.02 |
| *17* | | A | *R* | $R^{16}$ | 0.77±0.21 |
| *18* | | A | *R* | $R^{17}$ | 0.38±0.22 |
| *19* | *cis*-hydroxy-(*R*)-PZQ | A | *R* | $R^{18}$ | 5.39±1.48 |
| *20* | *cis*-hydroxy-(*S*)-PZQ | A | *S* | $R^{18}$ | inactive |
| *21* | *trans*-hydroxy-(*R*)-PZQ | A | *R* | $R^{19}$ | 5.59±1.39 |
| *22* | *trans*-hydroxy-(*S*)-PZQ | A | *S* | $R^{19}$ | inactive |
| *23* | | A | *R* | $R^{20}$ | 3.7±0.46 |
| *24* | | A | *R* | $R^{21}$ | 9.83±0.15 |
| *25* | | A | *R* | $R^{22}$ | inactive |
| *26* | | A | *R* | $R^{23}$ | inactive |
| *27* | | A | *S* | $R^{23}$ | inactive |
| *28* | | A | *R* | $R^{24}$ | inactive |
| *29* | | A | *R* | $R^{25}$ | inactive |
| *30* | | B | *R* | $R^1$ | 0.17±0.03 |
| *31* | | B | *R* | $R^4$ | 2.6±0.87 |
| *32* | | B | racemic | $R^{20}$ | 8.48±2.6 |
| *33* | | C | NA | $R^1$ | 8.67±1.74 |
| *34* | | D | racemic | $R^1$ | inactive |
| *35* | epsiprantel | E | racemic | $R^1$ | 0.41±0.27 |
| *36* | | E | racemic | $R^4$ | 1.27±0.55 |
| *37* | | F | racemic | $R^1$ | inactive |
| *38* | | F | racemic | $R^{26}$ | inactive |
| *39* | | F | racemic | $R^{27}$ | inactive |
| *40* | | G | NA | $R^1$ | inactive |
| *41* | | H | NA | $R^1$ | inactive |
| *42* | | I | NA | - | inactive |
| *43* | | J | NA | NA | inactive |

**Table S6. Activity of analogues against *Fh*.TRPM$_{PZQ}$(Thr$^{1270}$→Asn).** List of 43 analogues ordered by core structure ('A' through J') and substituents ($R^1$ through $R^{27}$) as shown in Figure S1. $EC_{50}$s for activation of *Fh*.TRPM$_{PZQ}$(Thr$^{1270}$→Asn) are listed, and analogues displaying either no measurable activity or activity ≤25% of PZQ at 50µM are categorized as 'inactive'.

**Movie S1. Location of (*R*)-PZQ binding site in *Sm*.TRPM$_{PZQ}$.** (*R*)-PZQ docking pose at base of VSLD (transmembrane helices S1-S4) displayed through a complete (360°) rotation of the *Sm*.TRPM$_{PZQ}$ homology model.
